# Centromeres are hotspots of cytosine methylation epimutations in a filamentous fungus

**DOI:** 10.64898/2026.03.03.709258

**Authors:** Mariana Villalba de la Peña, Clayton Hull-Crew, Taylor R. Hutter, Cynthia A. Vino, Peter Sarkies, Maria Colomé-Tatché, Frank Johannes, Andrew D. Klocko, Ilkka Kronholm

## Abstract

Epimutations are changes in chromatin modifications, such as DNA methylation or histone modifications. Some of these epigenetic changes can be inherited for several generations, and they potentially contribute to evolutionary processes. Estimates of epimutation rates now exists in a few species, but the presence and function of epigenetic marks are not conserved across different species. To understand the properties of epimutations in fungi, we performed a mutation accumulation experiment with the filamentous fungus *Neurospora crassa* and investigated spontaneous changes in DNA methylation and trimethylation of lysine 9 on histone H3 (H3K9me3) in the mutation accumulation lines. We observed that centromeric regions are hotspots of spontaneous DNA methylation changes in *N. crassa*. In these hotspot regions, DNA methylation changes were transmitted across mitoses, but changes occurring in euchromatin were not maintained. The rate of DNA methylation changes was around 30 000 fold faster than the genetic mutation rate. We did not observe spontaneous changes in H3K9me3 that were transmitted across mitoses. Our results show that while spontaneous epimutations occur in this species, they occur predominantly in gene poor heterochromatic regions, so their impact for evolutionary adaptation may be limited.

## Introduction

Epigenetic changes are inherited chromatin modifications that have the potential to regulate genome function without changing the underlying DNA sequence (Allis et al., 2009). A change in an epigenetic modification pattern that is inherited is referred to as an epimutation. Two of the most extensively studied epimutation types are cytosine methylation and histone post-translational modifications (PTMs).

Cytosine methylation is a key epigenetic modification that can silence transcription of genes, transposable elements (TEs), or other repetitive sequences (Mattei et al., 2022). The most common methylation site on DNA occurs when a methyl group is added to the fifth carbon within the pyrimidine ring of cytosine, usually in symmetric sequence context across both DNA strands, such as a CG motif, which allows methylation to be progated during DNA replication (Mattei et al., 2022). In addition CHG and CHH motifs can also be methylated.

In contrast, PTMs are chemical substituent groups that are covalently added to the N-terminal tails of histone proteins. These modifications can alter how chromatin is packaged in the nucleus and can mediate the silencing of certain genomic regions (Bannister and Kouzarides, 2011). There is evidence for inheritance of trimethylation of lysine 9 on histone H3 (H3K9me3) and trimethylation of lysine 27 on histone H3 (H3K27me3) (Moazed, 2011; Reinberg and Vales, 2018; Woodhouse and Ashe, 2020).

Moreover, DNA methylation can be randomly gained or lost, creating spontaneous epimutations (Becker et al., 2011; Schmitz et al., 2011; Johannes and Schmitz, 2019). DNA methylation epimutations occur at much higher rates than genetic mutations, yet these epimutations have high reversion rates (van der Graaf et al., 2015). Additionally, epimutations can occur in hotspots, which can be unstable in nematodes (Beltran et al., 2020), but are stable in plants (Hazarika et al., 2022). However, most studies to date have focused on single cytosine level changes, despite that spontaneous methylation changes across longer regions also occur (Denkena et al., 2021), and these changes maybe more functionally relevant than single site changes (Wibowo et al., 2016).

Understanding the biological significance of spontaneous epimutations is crucial, particularly in evolution, as they represent an additional layer of heritable variation (Johannes et al., 2009; Cortijo et al., 2014). Therefore, it is essential to characterize the properties of epimutations, including measuring their rates of gain and loss, identifying their genomic distribution, and assessing their heritability across generations. These insights are necessary to integrate spontaneous epimutations into evolutionary models (Kronholm and Collins, 2016). Moreover, available data on epimutations is often heavily skewed to a few model organisms, as most epimutation studies have been performed in plants or nematodes. However, it has been established that epimutations occur in many other organisms (Roquis et al., 2016; Calo et al., 2014; Torres-Garcia et al., 2020), necessitating a broader taxonomic perspective.

To understand the properties of epimutations, we studied the changes to cytosine methylation and H3K9me3 in the filamentous fungus *Neurospora crassa*. Neurospora has rapid generation time, and a small, well-characterized genome that contains many of the same epigenetic modifications conserved in higher eukaryotes, including cytosine methylation (Selker et al., 2003), H3K9me3 to demarcate permanently silent, gene poor and A:T rich constitutive heterochromatin (Lewis et al., 2009), and the H3K27me3 that denotes temporarily silent, gene rich facultative heterochromatin (Jamieson et al., 2013). The current model of cytosine methylation in *N. crassa* is that H3K9me3 is deposited on histones associated with heterochromatin. HP1 binds this mark and recruits the DNA methyltransferase DIM-2, which is thought to be solely responsible for all DNA methylation in Neurospora vegetative mycelium (Kouzminova and Selker, 2001; Tamaru et al., 2003; Freitag et al., 2004). Furthermore, DIM-2 has been shown to have activity to both unmethylated and hemimethylated sites *in vitro* (Shao et al., 2024), suggesting it may have both maintenance and *de novo* methylation activities. Interestingly, DIM-2 has no methyltransferase activity in the absence of HP1 or H3K9me3 (Shao et al., 2024), but when it is active, DIM-2-dependent cytosine methylation is not limited to the CG sequence context, but can methylate cytosines in CHG and CHH contexts (Shao et al., 2024).

Additionally, *N. crassa* has a unique genome defense system called repeat-induced point mutation (RIP), which detects duplicated genomic sequences during premeiotic stages and induces C *→* T transitions in these sequences (Selker, 1990). Because RIP introduces distinct mutations into each copy, large arrays of repeated sequences become increasingly divergent over evolutionary time (Cambareri et al., 1991). As a result, the genome of *N. crassa* contains few perfect repeats and it is possible to map short sequencing reads to centromeric regions, which is not possible for many other organisms (Smith et al., 2011).

We leveraged our previously generated mutation accumulation lines (Villalba de la Peña et al., 2023), to investigate spontaneous changes in DNA methylation and H3K9me3. Specifically, we set out to determine whether spontaneous epimutations occur in *N. crassa*; are epimutations inherited across asexual cell divisions; where in the genome epimutations occur; and what are the epimutation rates? Furthermore, since cytosine methylation depends on H3K9me3 in *N. crassa* (Tamaru et al., 2003; Shao et al., 2024), we assessed if methylation changes occur in the Neurospora genome due to alterations in underlying H3K9me3 or whether H3K9me3 remains constant while methylation changes.

## Results

### Spontaneous DNA methylation changes

We previously performed a mutation accumulation (MA) experiment with *N. crassa* (Villalba de la Peña et al., 2023), where ancestors were split into multiple MA lines, which were propagated by single spore descent by isolating a single conidia (asexual spore) and allowing the culture to grow and conidiate. Each MA line was transferred 40 times, which represents *≈* 1015 mitoses (Villalba de la Peña et al., 2023). Bottlenecks of single spore at each transfer minimize the strength of natural selection, which allows mutations to accumulate in each independent MA line (Halligan and Keightley, 2009). To address whether cytosine methylation epimutations would accumulate in the MA lines, we sampled ten MA lines from two independent MA pedigrees and performed bisulfite sequencing for samples taken after 5, 20, and 40 transfers (Figure 1A). We sequenced three biological replicates for both ancestors, i.e. from independent cultures and DNA extractions, two replicates at each time point for line 10, and one replicate per time point for rest of the MA lines, for 69 WGBS samples in total (Table S1).

**Figure 1:**
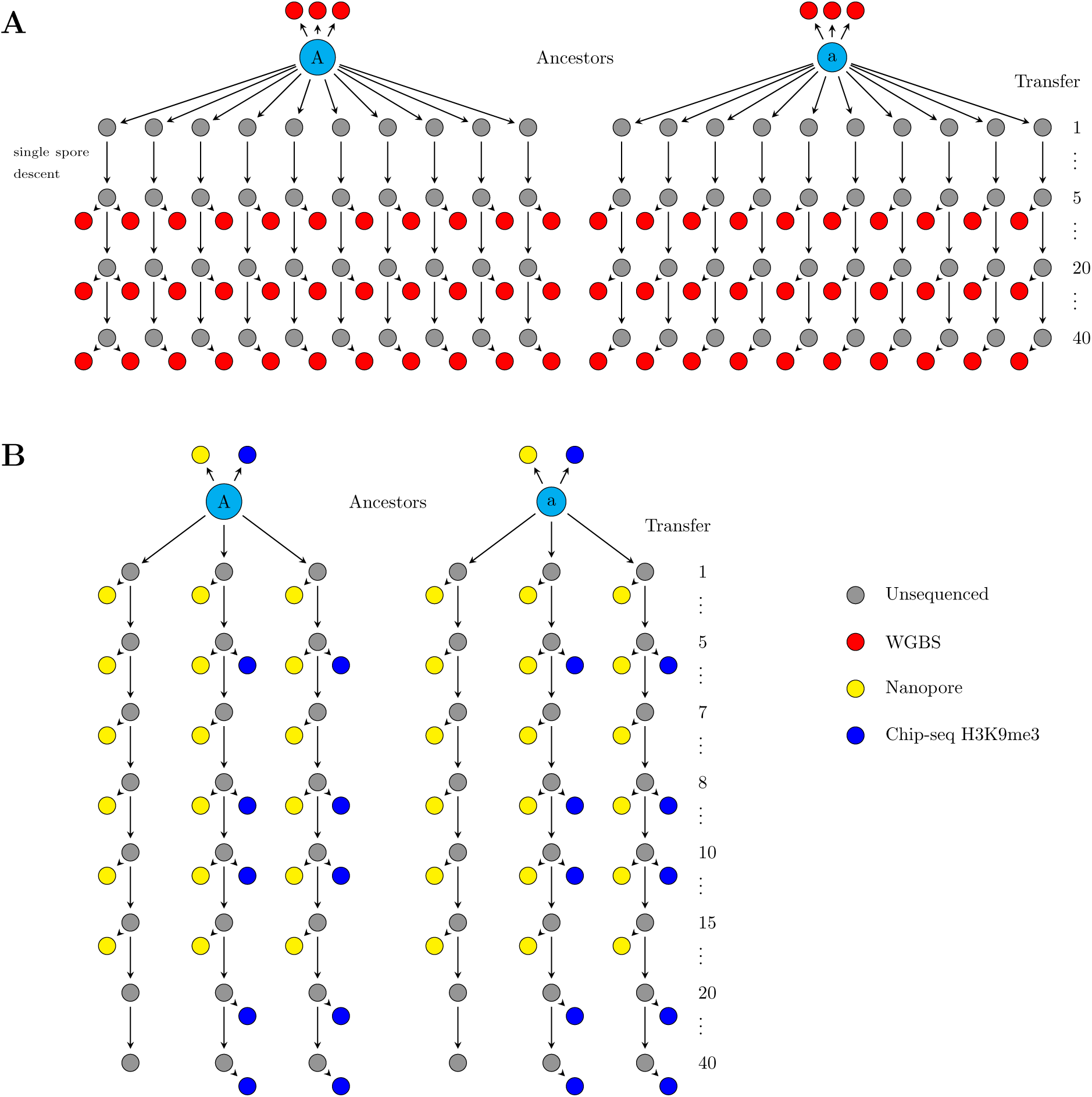
Samples taken from the two MA pedigrees for different experiments. A) Ten MA lines were sampled from the two pedigrees. Samples for sequencing were taken after 5, 20, and 40 transfers and DNA methylation was detected using bisulfite sequencing (red circles). B) Three additional MA lines were sampled from both pedigrees and samples were taken after 1, 5, 7, 8, 10, and 15 transfers for Nanopore sequencing and after 5, 8, 10, 20, and 40 transfers for H3K9me3 ChIP-seq.

First we examined where DNA methylation occurs in the MA ancestors, and observed that DNA methylation occurs mainly in centromeres and interspersed heterochromatic regions marked by H3K9me3 along the chromosomes (Figure S1 and S2). Euchromatin has very low levels of DNA methylation, and methylation is absent in genes (Figure S2). These observations are consistent with previous results about DNA methylation in *N. crassa* (Selker et al., 2003; Hosseini et al., 2020).

If spontaneous DNA methylation changes occur in the MA lines, and these changes are transmitted across mitoses, we should observe that the DNA methylation patterns of the MA lines diverge from each other. We calculated methylation divergence for CG, CHG, or CHH sequence context for each pair of MA samples and for each MA sample and its ancestor within a pedigree and plotted divergence against the number of mitoses that separate the samples (Figure 2). We observed that for single cytosines, DNA methylation divergence initially increased for the first 400 mitoses but leveled off and became saturated while variance increased substantially after 500 mitoses (Figure 2A). We tested whether this increase in methylation divergence over mitoses was significant by fitting a model of neutral epimutation accumulation (Shahryary et al., 2020) to the data. Compared to a null model of no accumulation, the neutral model was preferred in all contexts: *p* = 1.22 *×* 10*^−^*^8^, *p* = 4.18 *×* 10*^−^*^11^, and *p* = 7.16 *×* 10*^−^*^13^ for CG, CHG, and CHH respectively. However, the model of epimutation accumulation does not seem to capture the saturation effect in the data (Figure 2). We examined if specific samples cause many of the pairwise comparisons to have extreme divergence values, and we did identify four of such samples (Figure S3, supplementary results). We fitted a neutral accumulation model to the data with the outlier samples removed (Figure S4) and this improved the fit to the bulk of the data, but did not still capture the saturation effect (Figure 2A). We also tested fitting a purely phenomenological logistic growth model for the divergence and compared this to a null model of no divergence. The model with divergence was preferred in all sequence contexts, for CG context *p* = 2.12 *×* 10*^−^*^12^, for CHG *p* = 3.00 *×* 10*^−^*^14^, and for CHH *p* = 4.46 *×* 10*^−^*^16^. We cannot estimate epimutation rates using the logistic model, but this test shows that divergence among the MA lines increases over time, so cytosine methylation changes must be transmitted across mitoses to some extent.

**Figure 2:**
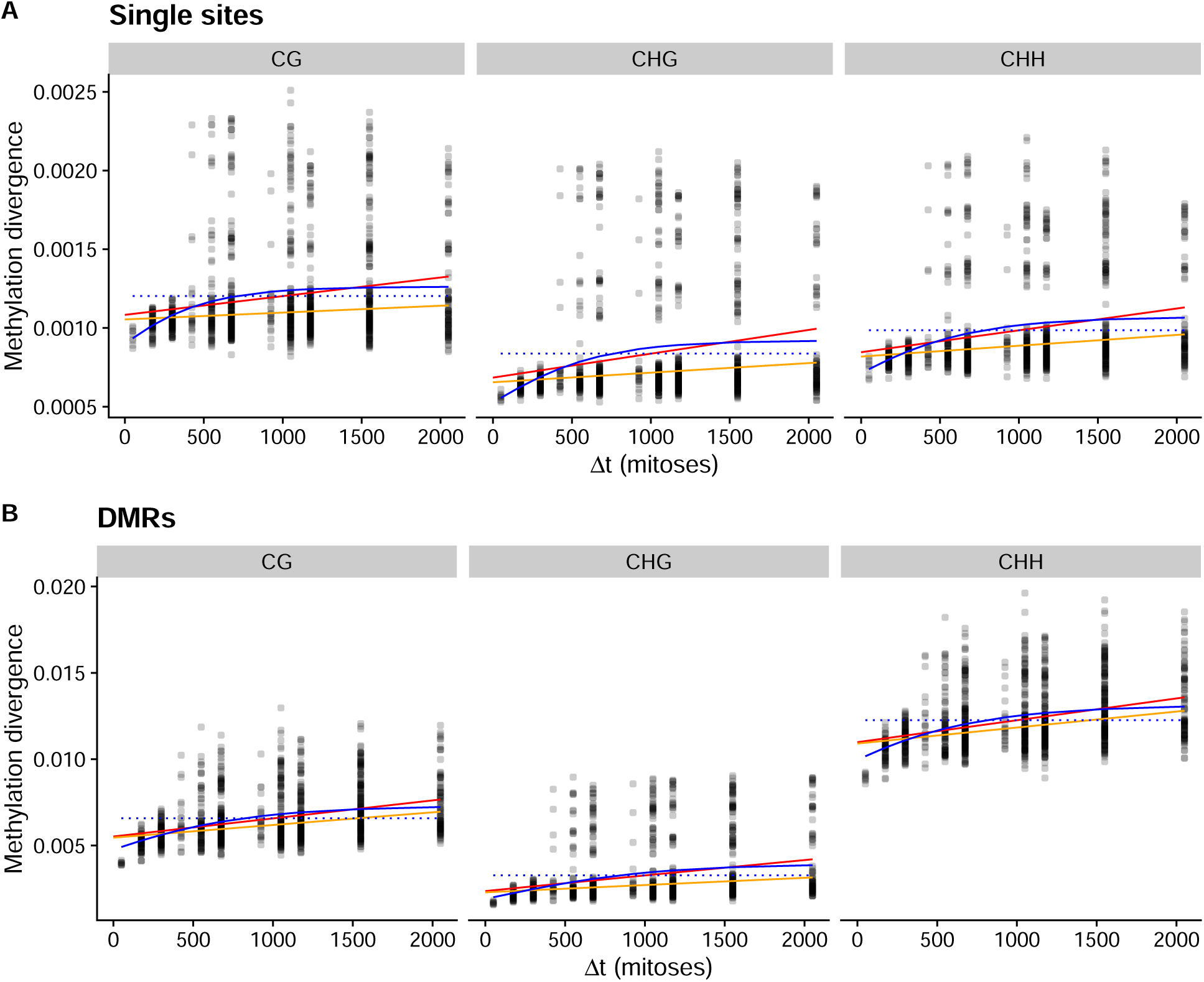
Pairwise divergence in cytosine methylation patterns among the MA lines and their ancestors. X-axis shows the number of mitoses separating the two lines that are being compared, and the y-axis shows cytosine methylation divergence. Red line shows a neutral epimutation accumulation model fitted to all data, orange line shows a neutral epimutation accumulation model fitted to data with outlier samples removed, and solid blue line shows a logistic model fitted to all data, and the dotted line shows null model of no increase in divergence. A) The divergence calculated from single cytosines, for CG, CHG, and CHH sequence motifs. B) The divergence calculated for DMRs in different sequence contexts, as in panel A.

We then tested whether differentially methylated regions (DMRs), defined as a genomic region containing five or more single cytosine methylation sites where methylation patterns change together, would also show divergence. We segmented the genome into 100 bp windows (bins) and determined which windows showed differences in methylation patterns among the samples, and those windows were classified as DMRs. Then we calculated methylation divergence for each pair of samples using the DMRs. We observed that the DMR divergence was more apparent for the first 400 mitoses than for single sites, especially in the CHH context (Figure 2B). We fitted a neutral epimutation accumulation model and compared it to a null model of no divergence, we observed that neutral accumulation was the preferred model in all contexts: *p <* 2.2 *×* 10*^−^*^16^ for all. This was also true for the data with outlier samples removed (supplementary results), and the logistic growth model was also preferred over the null in all contexts (*p <* 2.2 *×* 10*^−^*^16^ for all). As for single cytosines, the epimutation models did not capture the saturation effect in the DMR data (Figure 2B).

While our results show that methylation patterns must be transmitted over mitoses in *N. crassa*, the signal of divergence was not as clear as in *Arabidopsis thaliana* (van der Graaf et al., 2015), where divergence is approximately linear over many generations. To get a clearer picture of divergence in *N. crassa*, we investigated the genomic locations of the DMRs. When we plotted the DMR locations observed across MA lines on the *N. crassa* chromosomes, we observed that the vast majority of the DMRs occurred in centromeric and pericentromeric regions of each chromosome (Figure 3A and B). However, a minority of DMRs were also observed in either interspersed constitutive heterochromatic regions marked by H3K9me3, regions of facultative heterochromatin marked by H3K27me3, or in gene-rich euchromatin (Figure 3B).

**Figure 3:**
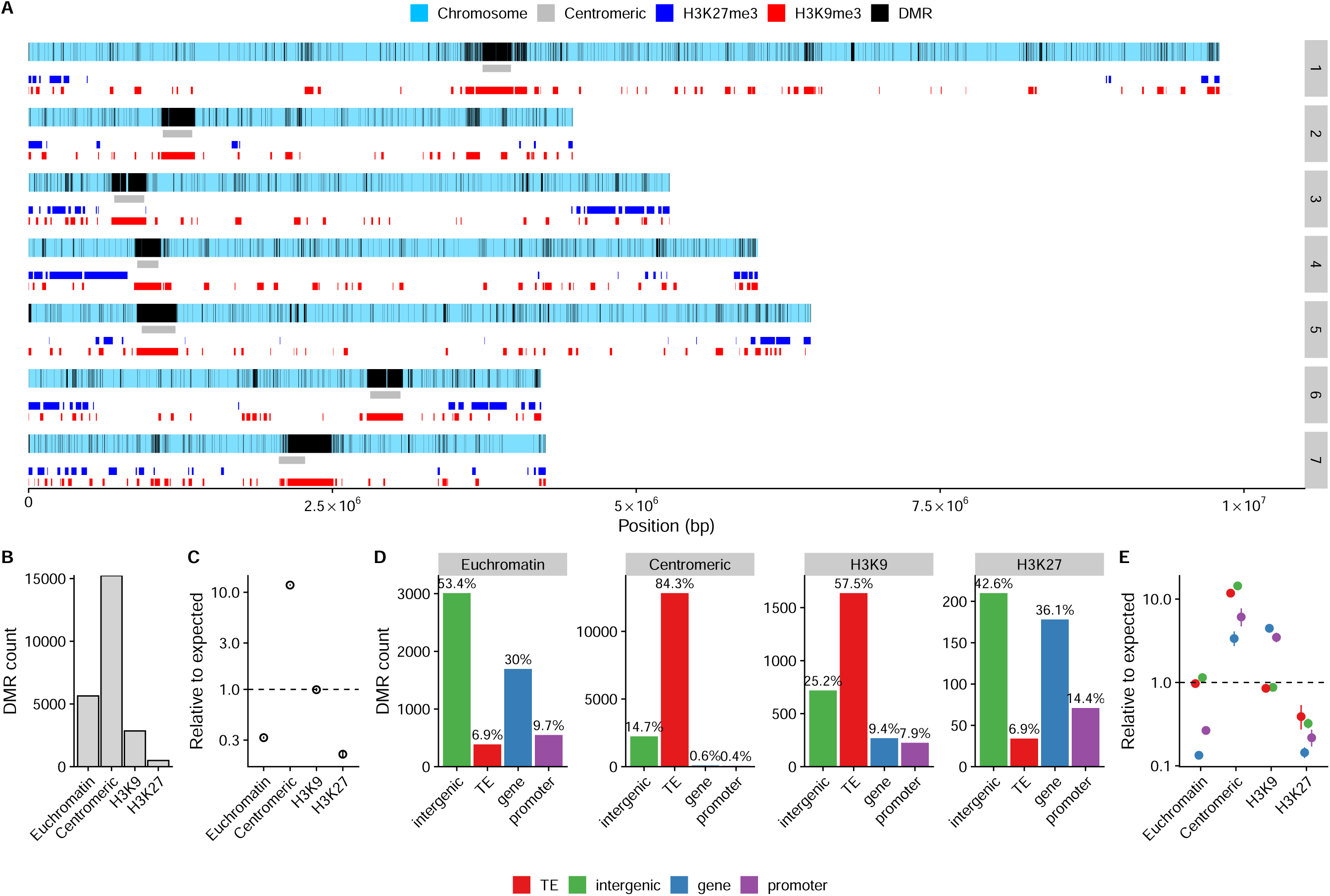
A) Locations of DMRs in the seven chromosomes of *N. crassa*, black vertical lines show DMRs observed across all MA lines. Colored boxes show locations of different chromatin domains. B) Counts of DMRs in the different domains. C) Model estimates for DMR rates in different domains, compared to random expectation. D) DMR counts and proportions split by the genome annotations for each chromatin domain. E) Model estimates for DMR rates for different annotations. In C) and E), the points show the estimates and lines indicate the 95% confidence intervals; most of them are obstructed by the points. Note the logarithmic scale for y-axis.

We tested if DMRs occurred randomly across the genome in the different regions (e.g. centromeric, or euchromatin etc.) by fitting a model of DMR counts in each region, accounting for region size. We observed that DMRs were tenfold more common in centromeric regions compared to random expectation (*p <* 2 *×* 10*^−^*^16^), while DMRs were rarer than expected in euchromatic regions, with only 32% DMRs observed relative to expected (*p <* 2 *×* 10*^−^*^16^). DMRs were also underrepresented in H3K27me3 regions, only 21% of expected DMRs were observed (*p <* 2*×*10*^−^*^16^). For regions marked by H3K9me3 outside the centromeric regions, there was no difference with the expected amount of DMRs, *p* = 0.641 (Figure 3C). Furthermore, we tested whether the enrichment of DMRs in the centromeric regions was robust to DMR specification, and observed that DMRs were enriched in the centromeric regions for all tested window sizes (Figure S5A, and supplementary results).

Transposable elements are often associated with DNA methylation and DMRs, so we checked if DMRs overlapped any particular TE annotations to examine if the presence of TEs could promote higher DMR occurrence in centromeres. In centromeric regions the majority of DMRs are found in sequences annotated as transposable elements (Figure 3D), while in euchromatic regions and regions marked by H3K27me3, the proportion of TE annotations was much lower, with DMRs occurring in both genic and promoter sequences. For interspersed (non-centromeric) constitutive heterochromatic regions marked by H3K9me3, DMRs occur mainly in TEs, but also in genic sequences (Figure 3D).

To test whether the observed distribution of DMRs could be explained by random placement with respect to genomic annotations, we fit a model that accounts for the effects of the chromatin domain, sequence annotation, and their interaction. DMRs were significantly enriched in TEs and other genomic features in centromeric regions (*p <* 2 *×* 10*^−^*^16^), whereas in euchromatic regions, DMRs occurred in TEs at levels consistent with random expectation (*p* = 0.544; Figure 3E). In H3K9me3-marked regions DMRs were slightly depleted in TEs, only 85% of DMRs that would be expected by chance occurred in TEs (*p* = 5.47 *×* 10*^−^*^11^; Figure 3E). Furthermore, in H3K27me3-marked regions, DMRs we depleted in TEs, only 39% of DMRs that would be expected chance occurred in TEs (*p* = 3.88 *×* 10*^−^*^8^; Figure 3E). We also observed that DMRs were more often absent from genes and promoters in facultative heterochromatic regions marked by H3K27me3 or in euchromatic genes (Figure 3E). In contrast, predicted genes and their promoters in centromeric or interspersed H3K9me3 regions are enriched for DMRs (Figure 3E).

Our observation that DMRs occur in TEs and intergenic sequences at very similar rates in all regions (Figure 3E), prompted us to investigate whether differences in TE composition between centromeric and non-centromeric regions could explain the overabundance of centromeric DMRs, but we found no evidence explaining the observation (Supplemental results, and Figure S6). Therefore, we concluded that it is not specifically TE sequences that promote additional centromeric DMRs, but additional centromeric factors must explain increased rate of DMRs in centromeres or their retention.

The uneven distribution of DMRs across the genome prompted us to investigate whether different genomic regions have altered methylation pattern divergence. We plotted divergence in DMRs for different regions of the genome separately, and observed that the MA lines diverged only for centromeric DMRs (Figure S7), with no divergence in other genomic regions (Figure S7). The same pattern could be observed for methylation divergence at single sites (Figure S8). Since divergence happens only in centromeric regions, which represent a small proportion of the whole genome, and if epimutations rates are high, this could potentially cause the observed saturation effect. To both confirm that divergence is happening using an orthogonal technique, and to obtain a more accurate divergence rate estimate using denser sampling of the MA pedigree, we sampled six additional MA lines from the two pedigrees, with six time points per line closer to each other in time than the three time points in WGBS data, and performed long-read Nanopore whole genome sequencing (Figure 1B, Table S2). These data confirmed that divergence in methylation patterns happened exclusively within centromeric DMRs (Figure S9) and single sites (Figure S10) with no divergence observed for other genomic loci. This denser sampling of the MA pedigree showed that the initial divergence of centromeric cytosine methylation increases linearly over the shorter time period captured within the Nanopore-seq dataset, before reaching saturation (Figure 4). Divergence in methylation patterns is caused by both gain and loss of methylation events, and cannot be explained centromeric regions just losing methylation gradually (Figure S11 and supplementary results). Divergence for DMRs is also robust to parameter choices in the DMR segmentation (Figure S5B). Moreover, methylation divergence is not compatible with a simple model of nucleosome position drift (Figure S12 and supplementary results). Together, we conclude that DNA methylation changes are transmitted across mitoses solely within centromeric regions in *N. crassa*.

**Figure 4:**
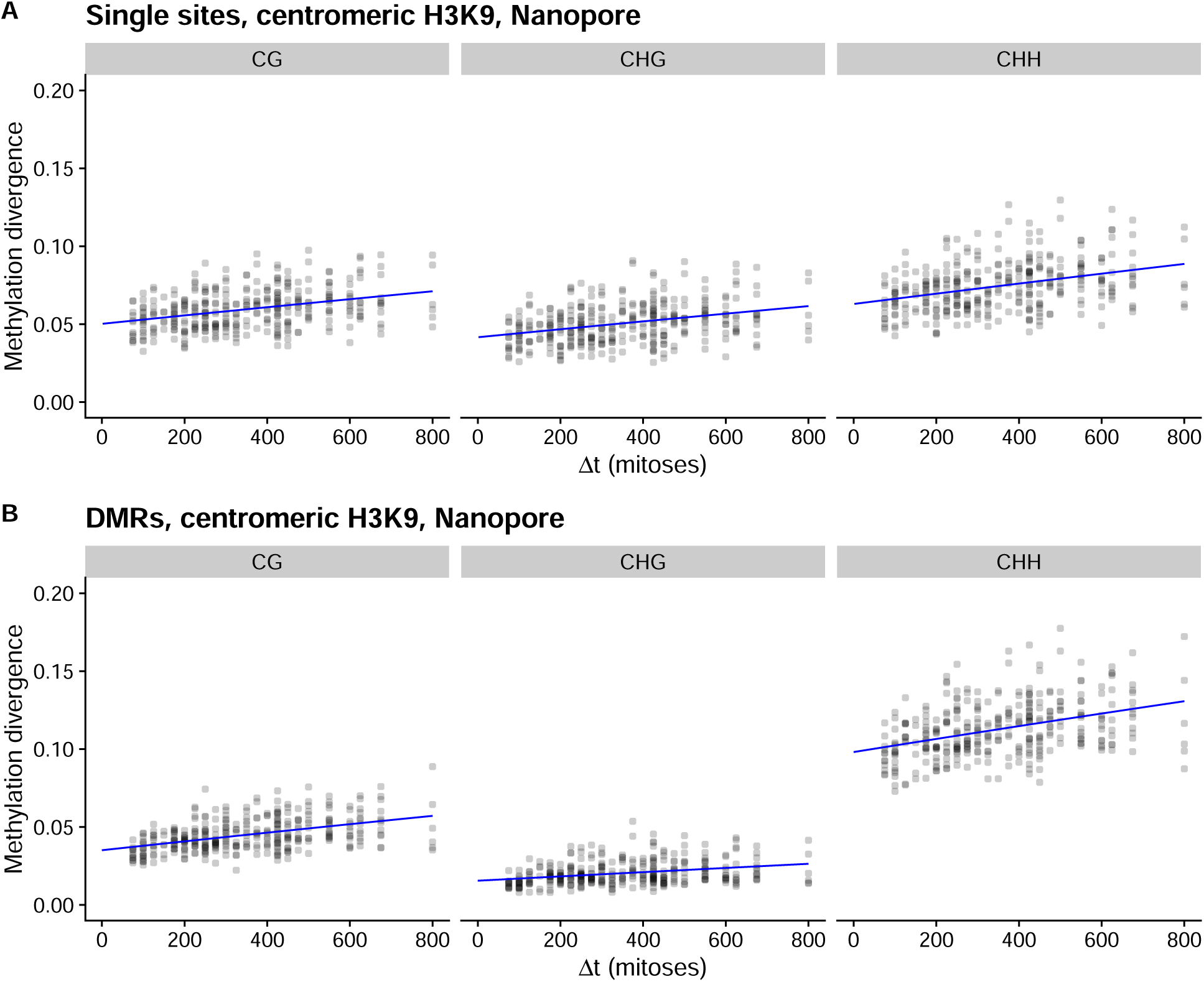
Pairwise divergence in methylation patterns between individual MA lines and their ancestors in centromeric regions for the Nanopore-seq dataset. The x-axis shows the number of mitoses separating the two compared strains, while the y-axis shows the cytosine methylation divergence. Blue line is a fit from the AlphaBeta neutral epimutation accumulation model. A) The divergence calculated for single sites. B) The divergence calculated for DMRs.

### Rate of spontaneous cytosine methylation changes

The linear increase of DNA methylation changes in the Nanopore-seq data allowed us to fit models of epimutation accumulation to these data, and to calculate the rate of spontaneous methylation changes in centromeric regions. For single cytosines, a model with neutral accumulation of epimutations was strongly preferred over the null model of no epimutation accumulation in CG (*p* = 1.09 *×* 10*^−^*^9^), CHG (*p* = 9.11 *×* 10*^−^*^9^), and CHH (*p* = 1.65 *×* 10*^−^*^9^) sequence contexts (Figure 4A). For DMRs a model of neutral accumulation of epimutations was also preferred over the null model in all sequence contexts, CG (*p <* 2.2 *×* 10*^−^*^16^), CHG (*p* = 3.25 *×* 10*^−^*^7^), and CHH (*p* = 3.01 *×* 10*^−^*^12^).

Spontaneous DNA methylation gains for individual cytosines occurred at a rate of 1.5 to 2.2 *×*10*^−^*^5^ per cytosine per mitosis for all sequence contexts, and the estimates were within the bounds of measurement uncertainty from each other (Table 1). Methylation losses occurred at rates that were around 3–5 times faster: around 7–8 *×*10*^−^*^5^ per cytosine per mitosis consistent for all sequence contexts (Table 1). Generally, rate estimates for DMRs were within the bounds of measurement uncertainty from the single site estimates, with the exception of rate of methylation gain for CHG context, which had a slightly higher rate (Table 1). There were also slight differences for the DMR rates in different sequence contexts (Table 1). Then we examined if we could obtain comparable estimates from the initial divergence of the WGBS data, and observed that estimates for single sites were consistent with estimates of the Nanopore dataset, but estimates for DMRs were not (see supplementary results and Figure S13). Even though the rate of loss of methylation per cytosine is higher, some methylation is expected to be maintained, since the expected number of cytosine methylation events increases as the number of unmethylated cytosines increases, leading eventually to a dynamic equilibrium. We calculated this equilibrium proportion of methylated cytosines based on Markov chain theory, which for these rates is around 16% for CG and CHG, and around 23% for CHH context.

**Table 1:** Estimates of rates of spontaneous gains (*α*), losses (*β*), and their ratio (*β/α*) for DNA methylation in centromeric regions. Units for the rates are: epimutations per cytosine per mitosis for single sites, and epimutations per bin per mitosis for DMRs. Square brackets show the 95% bootstrap interval for the estimate.

| Parameter | Context | DMRs | Single sites |
| --- | --- | --- | --- |
| $\alpha$ | CG | $1.72 [1.37, 2.05] \times 10^{-5}$ | $1.62 [1.18, 2.09] \times 10^{-5}$ |
| $\beta$ | CG | $8.79 [6.99, 10.50] \times 10^{-5}$ | $8.55 [6.25, 11.11] \times 10^{-5}$ |
| $\beta/\alpha$ | CG | $5.12 [5.11, 5.12]$ | $5.30 [5.29, 5.31]$ |
| $\alpha$ | CHG | $0.79 [0.53, 1.06] \times 10^{-5}$ | $1.54 [1.07, 2.06] \times 10^{-5}$ |
| $\beta$ | CHG | $5.66 [3.79, 7.58] \times 10^{-5}$ | $8.19 [5.70, 10.94] \times 10^{-5}$ |
| $\beta/\alpha$ | CHG | $7.14 [7.13, 7.15]$ | $5.31 [5.30, 5.33]$ |
| $\alpha$ | CHH | $3.01 [2.24, 3.75] \times 10^{-5}$ | $2.17 [1.59, 2.83] \times 10^{-5}$ |
| $\beta$ | CHH | $7.19 [5.35, 8.94] \times 10^{-5}$ | $7.18 [5.26, 9.35] \times 10^{-5}$ |
| $\beta/\alpha$ | CHH | $2.39 [2.38, 2.39]$ | $3.31 [3.30, 3.31]$ |

### Genetic mutations and DNA methylation changes

In principle, all of the observed DNA methylation changes could be governed by genetic mutations. To examine the relationship between genetic mutations and methylation changes, we compared patterns of genetic mutations and DNA methylation changes in our MA lines.

We previously determined the genetic mutations occurring in the MA lines, with each MA line having a median of 33 new genetic mutations by the 40th transfer (Villalba de la Peña et al., 2023). For the 20 MA lines sequenced by bisulfite sequencing, we observed a total of 725 genetic mutations at the end point of the MA experiment. Of these mutations, 301 were single nucleotide mutations where cytosine was either the ancestral or the new base. In total, we observed 1 115 460 cytosines with variable methylation patterns across all MA lines and time points, involving 5.68% of all genomic cytosines, comprising a total of 24 208 DMRs. Given the amount of cytosine methylation changes compared to genetic mutations, we can outright reject the explanation that genetic mutations control all cytosine methylation changes in *cis*.

We further assessed if genetic mutations were controlling DMRs in *cis*. We examined the intermediate MA time points from transfers 1, 5, 7, 8, 10 and 15 within the six MA lines L2, L5, L11, L23, L25, and L31 subjected to Nanopore-seq (Figure 1B). From these Nanopore-seq data, we tested if known genetic mutations were present or absent at the time points when the overlapping DMRs occurred. These MA lines accumulated 121 genetic mutations by transfer 40. First, considering genetic mutations that could potentially control DMRs in *cis*, only 14 of these observed mutations overlapped with DMRs in at least at one time point. If the genetic mutations control DMRs, we would expect any DNA methylation change to be stable from the time point when that genetic mutation occurred. Instead, we observed that DMRs did not necessarily occur at the same time as the overlapping genetic mutation; when DMRs and genetic mutations did overlap, the DMR had the potential to disappear in subsequent time points or change without an overlapping genetic mutation (Figure S14).

Next, we assessed if the data were consistent with genetic mutations controlling multiple DMR changes in *trans*. We examined all genetic mutations and all DMR changes that occurred between different transfers within each line. Across six transfers intervals for the six MA lines, we observed seven intervals where no genetic mutations occured in that MA line. Yet, in those cases, a typical number of DMRs changes was observed in those lines (Figure 5). DMR changes without genetic mutations also happened in both centromeric regions and the rest of the genome (Figure 5). The pattern was similar for methylation changes at individual cytosines (Figure S15). Furthermore, DMR changes do not significantly increase in intervals with genetic mutations (Figure S16), which would be expected if genetic changes profoundly affect methylation patterns. Thus, the majority of methylation changes are not under genetic control and instead represent spontaneous changes, because they arise without co-occurring genetic mutations. While we cannot exclude the possibility that an isolated genetic mutation in an individual MA line may affect methylation patterns, we conclude that genetic mutations cannot explain the broad pattern of cytosine methylation divergence in centromeres.

**Figure 5:**
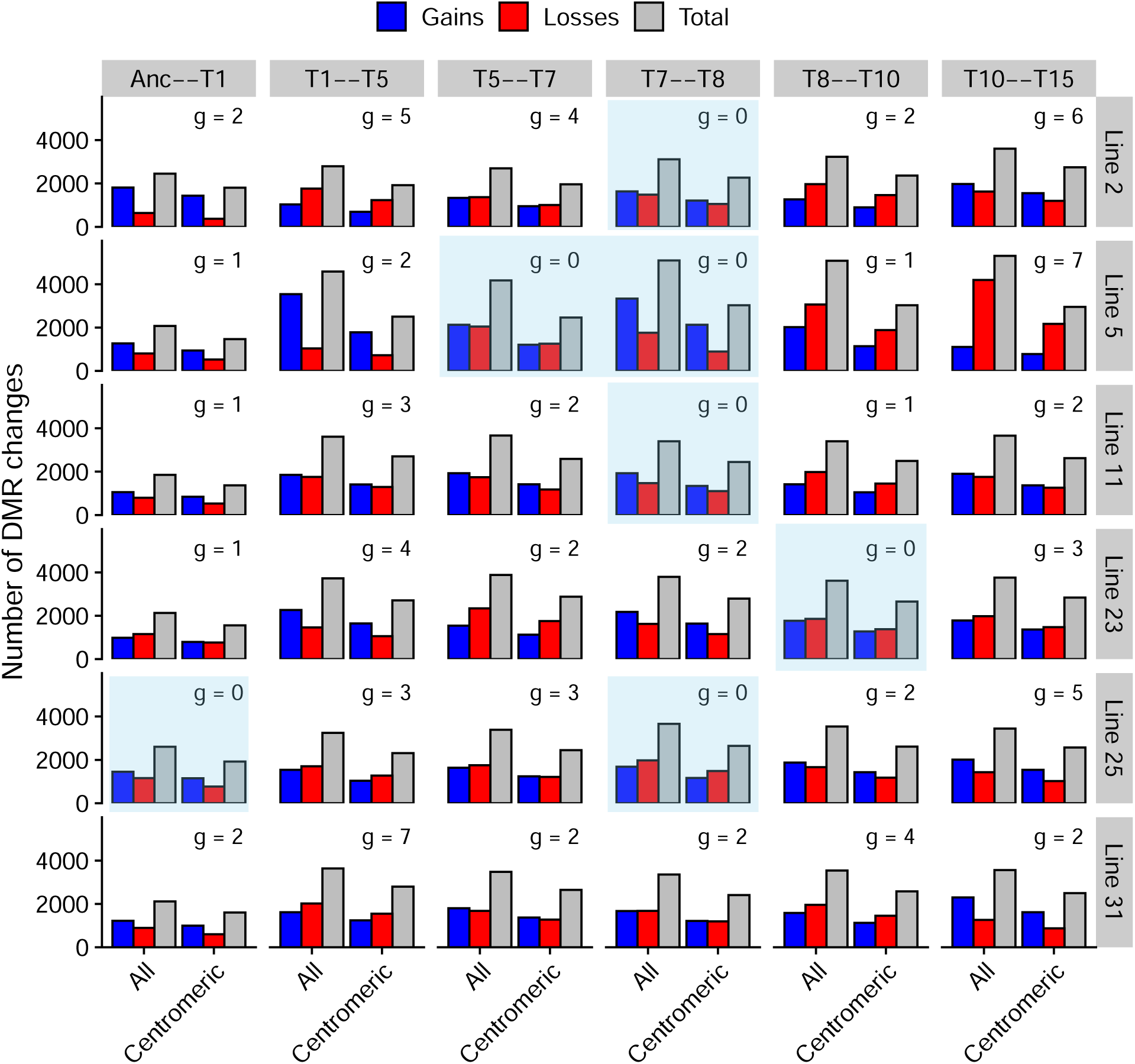
Number of DMR changes for the whole genome and centromeric regions across the transfer intervals (columns) for all MA lines (rows) sequenced with Nanopore-seq. Number of genetic mutations (g) occurring in each interval is shown on top of the bars, and intervals where no genetic mutations occurred are highlighted in blue.

### DNA methylation and H3K9me3

In *Neurospora crassa*, DNA methylation is primarily found in heterochromatic regions marked by H3K9me3, as H3K9me3 is bound by Heterochromatin Protein 1 (HP1) for recruiting the DNA methyltransferase DIM-2, which is thought to be responsible for all DNA methylation in Neurospora (Tamaru et al., 2003; Freitag et al., 2004; Honda and Selker, 2008). To investigate whether the observed DMRs exclusively represent DNA methylation changes, or whether the underlying H3K9me3 enrichment is altered, we performed H3K9me3 Chromatin Immunoprecipitation-sequencing (ChIP-seq) on four MA lines (L5, L11, L25, and L31) from transfers 5, 8, 10, 20, and 40 (Figure 1B and Table S3).

We assessed the overlap between DMRs and H3K9me3 regions in the MA ancestors and in MA lines from transfers 5, 8, and 10 for which we had both DNA methylation and H3K9me3 data. In these samples, we identified 14 503 unique DMRs, 90.64% of which overlapped with stable H3K9me3-enriched regions. Therefore, the vast majority of DMRs occur without accompanying changes to H3K9me3, and DNA methylation can be altered independently despite H3K9me3 being stable. Interestingly, of the DMRs that did not overlap H3K9me3, 95.1% were associated with gains of cytosine methylation (Figure S17A). DMRs that overlapped with H3K9me3 occurred mainly in TEs and intergenic sequences, while non-overlapping DMRs occurred much more frequently in euchromatic genes and promoters (Figure S17B).

While DMRs outside centromeric H3K9me3 were not transmitted across mitoses, to further understand the formation of these transient euchromatic DMRs, we investigated whether spreading of DNA methylation from nearby TEs could explain our observations. Using the bisulphite sequencing dataset, we picked the DMRs that occurred in euchromatic genes and promoters, and checked the distance to the nearest TE. Then we compared the distribution of obtained distances, to distancas of randomly picked genes and promoters, and observed that there was no difference (Supplementary Figure S18). Thus these euchromatic DMRs were not closer to TEs than random genes and promoters, suggesting that methylation spreading from TEs is unlikely to explain the formation of these transient DMRs.

### Spontaneous changes in H3K9me3

After observing that DNA methylation changes are transmitted across mitoses in centromeres, we investigated whether H3K9me3 patterns also change. As expected, we found that H3K9me3 is enriched in the MA lines in regions previously characterized as heterochromatic: centromeres, telomeric regions, and interspersed heterochromatic regions (Lewis et al., 2009; Jamieson et al., 2016; Klocko et al., 2019). We called regions of H3K9me3 by two different methods that gave concordant results, and observed that generally larger regions of H3K9me3 are stable, without substantial differences in enrichment, in the MA lines (Figure S19, S20, and S21). Further, the presence or absence of H3K9me3 peaks is highly correlated between MA lines and transfers (Figure S22), highlighting how larger regions of H3K9me3 in the Neurospora genome were stable during the MA experiment.

When then performed a fine scale analysis for detecting small scale differences in H3K9me3. First we looked for the presence and absence of smaller peaks. In total, we identified 271 variable peaks in the MA lines. We calculated the divergence in the presence or absence of these H3K9me3 peaks among the MA lines. However, the H3K9me3 peak divergence did not concordantly increase over time in the MA pedigree (Figure S23), indicating that these differences in small peaks of H3K9me3 are not stably transmitted across mitoses.

Peak-calling software may lack sensitivity and resolution to detect subtle changes, and peak boundaries can be ambiguous. We therefore conducted a complementary analysis by dividing the whole genome and centromeric regions into 5000 bp and 500 bp bins, respectively, and calculated the divergence in H3K9me3 read counts across bins and time points. H3K9me3 divergence did not increase across more distant MA lines within the MA pedigree (Figure S24, and supplementary results). This pattern was consistent across the whole genome, in the centromeric regions, or when centromeric regions were excluded (Figure S24). In addition, H3K9me3 regions do not undergo significant expansions or contractions over time (Figure S25, and S26). Together, while H3K9me3 can have slight variations in enrichment, we did not find any evidence that changes in H3K9me3 patterns are transmitted across mitoses. The subtle variation in H3K9me3 patterns may represent technical differences in ChIP-seq experiments rather than a biological phenomenon. Overall, H3K9me3 regions appear to be stable in the MA lines.

## Discussion

### Spontaneous cytosine methylation changes

We observed spontaneous cytosine methylation changes in the MA lines. Our results are broadly consistent with known cytosine methylation activity of DIM-2. Majority of DNA methylation was observed in heterochromatic regions marked by H3K9me3, which is expected as DIM-2 requires H3K9me3 and HP1 for activity (Freitag et al., 2004; Shao et al., 2024). Moreover, we observed changes in methylation patterns in CG, CHG, and CHH contexts consistent with known DIM-2 activity. However, the *N. crassa* genome encodes another putative DNA methyltransferase, RID, which may be responsible for cytosine methylation during RIP (Freitag et al., 2002). Hypothetically, during RIP, cytosine methylation and subsequent deamination functions within a C *→* T base editing pathway (Selker, 1990; Freitag et al., 2002). However, since RIP functions in meiosis (Selker, 1990) but the propagation of the MA lines was asexual, any RID-dependent methylation during RIP is unlikely to be responsible for the DNA methylation changes we observed, although it is possible that RID may have some, as of yet uncharacterized role outside RIP. Nevertheless, the relative contributions of DIM-2 and RID to the DNA methylation changes observed here, warrant future investigations.

The evolutionary origin of constitutive heterochromatin in *N. crassa* is likely to be the action of the RIP system. Over evolutionary time, RIP has degraded ancestral duplicated regions and TEs, and these regions have become AT rich and subsequently RIP mutated sequences are the initial step for heterochromatin formation allowing DNA methylation (Lewis et al., 2009; Rountree and Selker, 2010; Zemach et al., 2010; Hosseini et al., 2020). In *N. crassa* ancestral duplications affected by RIP and regions marked by H3K9me3 completely overlap (Lewis et al., 2009; Wang et al., 2020; Villalba de la Peña et al., 2023). Some of these AT rich sequences are likely to be consistently targeted by DIM-2 for methylation, as complementing the *dim-2* mutant restores methylation to heterochromatic regions (Kouzminova and Selker, 2001). However, our results show that not all cytosine methylation is constant in *N. crassa*, and some sites in heterochromatin can have variable methylation. We also observed that H3K9me3 was constant and did not change in heterochromatic regions, showing that cytosine methylation changes were not due to changes in underlying H3K9me3, but spontaneous changes in cytosine methylation happen over a constant H3K9me3 landscape. What are the sequence features in H3K9me3 marked heterochromatic regions that lead to constant methylation and which features allow for variability will need to be explored further.

### Inheritance of cytosine methylation changes

Strikingly, while spontaneous cytosine methylation changes occurred across the whole genome, the MA lines only diverged in their methylation patterns centromeric and pericentromeric regions. This divergence was not consistent with only variation in *de novo* targeting of methylation, but maintenance of methylation patterns across mitoses is required to explain the results. In contrast to plants and animals which have different enzymes for maintenance and *de novo* methylation, DIM-2 in *N. crassa* has both *de novo* and maintenance activities (Shao et al., 2024). In *in vitro* assays DIM-2 had higher activity to a hemimethylated CG site than to an unmethylated CG site (Shao et al., 2024), so maintenance methylation that allows methylation patterns to be transmitted across mitoses is consistent with known DIM-2 activity. However, as DNA methylation patterns were only transmitted across mitoses in centromeric and pericentromeric regions, these loci must harbor a unique signal that allows for the maintenance of DNA methylation patterns. Importantly, the presence of H3K9me3 cannot be this signal, as DNA methylation was transmitted independently of H3K9me3 changes, and DNA methylation changes were not transmitted across mitoses in interspersed heterochromatic regions even if these regions harbor H3K9me3.

One hypothesis is that DNA demethylation actively occurs outside centromeres but does not occur within centromeric DNA, perhaps because the centromere is inaccessible to DNA demethylation machinery. There are some reports that DNA demethylation is an active process in fungi (Antequera et al., 1985), but this is unclear in *N. crassa*. In mammals, DNA demethylation is catalyzed by TET proteins which oxidize 5-methylcytosine through several reactions to create an abasic site, which is then repaired by the base excision repair pathway to restore unmethylated cytosine (Wu and Zhang, 2014). In plants, DNA demethylation is catalyzed by ROS1, which is a bifunctional DNA glycosylase that cleaves 5-methylcytosine from DNA for DNA demethylation (Agius et al., 2006). *Neurospora* does not appear to encode any TET gene homologs, so if *N. crassa* has DNA demethylation activity, a DNA glycosylase may be responsible. Alternatively, DNA demethylation could be mediated by simple dilution of methylated cytosines during multiple DNA replications if hemimethylated sites present after DNA synthesis are not remethylated (Wu and Zhang, 2014). However, a dilution based mechanism implies that an active mechanism must maintain DNA methylation activity in centromeric regions, compared to other interspersed heterochromatic regions. In *N. crassa* centromeres are enriched for the histone H3 variant CenH3, as well as the centromere proteins CEN-C and CEN-T, and show extensive overlap with H3K9me3 (Smith et al., 2011). If present, an active cytosine methylation mechanism could employ the centromeric histone variant CenH3 (Smith et al., 2011), where either the deposition of CenH3, or a PTM on its disordered N-terminal tail, could signal for HP1/DIM-2 recruitment. Future experiments should elucidate the underlying mechanisms for the specific maintenance of cytosine methylation on (peri)centromeres.

### DNA methylation in euchromatin

Interestingly, we did a detect gains of euchromatic DNA methylation where H3K9me3 was not present, which appear inconsistent with the H3K9me3-dependent mechanism mediating DIM-2 methylation. Yet, it is also possible that our Chip-seq was not sensitive enough to detect H3K9me3 enrichment at these sites, if the change in chromatin state is very dynamic. We detected these euchromatic cytosine methylation gains using both bisulfite and nanopore sequencing, arguing against their presence due to technical artifacts, but these changes were not transmitted across mitoses in a stable manner. As such, it is unclear what these ephemeral changes represent, and whether they have any biological function.

DNA methylation in *N. crassa* euchromatin has been observed previously, as DIM-2-dependent DNA methylation has been detected in the promoter of the *N. crassa frq* gene, which requires the chromatin remodeler CHD1 (Belden et al., 2011). There are also observations that remodeling and deposition of H3K9me3 at the *frq* locus depends on the chromatin remodeler SAD-6 (Eugene Gladyshev, personal communication), which initiates transcriptional silencing in mitotic cells in *N. crassa* (Carlier et al., 2024). SAD-6 is the fungal ortholog of the human ATRX protein (Aguilera and López-Contreras, 2023), which mediates DNA methylation changes in repeated sequences in humans (Gibbons et al., 2000) and resolves G-quadruplex DNA complexes (Teng et al., 2021). Nucleosome depleted chromatin, caused by either aberrant protein binding (Carlier et al., 2024) or a G-quadruplex (Teng et al., 2021) can trigger silencing and heterochromatization. SAD-6 may deposit a H3K9me3-marked histone in perturbed regions (Aguilera and López-Contreras, 2023), for potential cytosine methylation in these perturbed sites. The stochastic nature of these euchromatic cytosine methylation events may even be explained by the random meeting of a replication fork with a G-quadruplex, which would be resolved by ATRX (Truch et al., 2022). It is unknown if chromatin remodeling by SAD-6 also causes DNA methylation in *N. crassa*, as previous work used DNA methylation deficient (Δ*dim-2*) strains (Carlier et al., 2024). So, whether our observations of euchromatic DNA methylation represent SAD-6 dependent H3K9me3 deposition needs to be investigated further.

### Rates of spontaneous cytosine methylation changes

Comparison of our *N. crassa* epimutation results to those from different species reveals remarkable divergence. MA studies in the plant *Arabidopsis thaliana*, revealed that DNA methylation epimutations are only transmitted across generations in the CG sequence context, with no appreciable divergence in individual cytosines or DMRs for CHH or CHG contexts (Becker et al., 2011; Schmitz et al., 2011; van der Graaf et al., 2015; Denkena et al., 2021). Furthermore, in *A. thaliana* epimutations primarily arise in genic and intergenic regions, and are strongly depleted in TEs (Denkena et al., 2021; Johannes and Schmitz, 2019), which contrasts the sequence context-independent epimutations occurring in TE-rich centromeric and heterochromatic regions in *N. crassa*. The differences between Neurospora and plants for cytosine sequence context can be explained by the cytosine specificity of DIM-2 and the plant maintenance DNA methyltransferase MET1, which prefers CG motifs (Shao et al., 2024). Further, the rate of spontaneous DNA methylation changes in both systems happens orders of magnitude faster than genetic mutations, and in both plants and fungi the rate of methylation loss is faster than the rate of methylation gain. In *A. thaliana* the rate of methylation gain in CG context is roughly 16 times faster, and the rate of loss 7 times faster than in *N. crassa* (van der Graaf et al., 2015). Importantly, while DNA methylation changes can contribute to *A. thaliana* phenotypic variation (Johannes et al., 2009; Cortijo et al., 2014), it appears that contribution to phenotypic variation may be limited in *N. crassa*, given that the inherited methylation changes occur only in gene-poor centromeric regions. Furthermore, we have no evidence that the centromeric DNA methylation changes in Neurospora compromise centromere function, as DNA sequencing did not reveal any changes in chromosome number in the MA lines (Villalba de la Peña et al., 2023), and loss of cytosine DNA methylation alone does not prevent normal centromere formation (Smith et al., 2011).

Overall, there is clear diversity in the underlying epigenetic mechanisms driving spontaneous epigenetic changes across eukaryotic taxa. It can be speculated that ecological factors may determine whether species have epigenetic systems that produce spontaneous epigenetic variation, which should become more evident as more epimutation studies accumulate. Even if the cytosine methylation changes that occur in Neurospora are neutral, they may have a practical use, as cytosine methylation changes have recently been used as epigenetic clocks to time branches of long lived forest trees (Hofmeister et al., 2020) or improve phylogenetic resolution on short time scales (Yao et al., 2023). Thus, neutral epigenetic variation has the potential to improve population genetic inference (Sellinger et al., 2024).

## Methods

### Mutation accumulation experiment

For details of the MA experiment, see Villalba de la Peña et al. (2023). Briefly, two ancestors of the FGSC 2489 (74-OR23-1VA) wild type background that are isogenic apart from their mating types were divided into 20 lines each, 40 lines total. Lines were transferred by plating asexual spores, picking a single colony to a test tube to produce asexual spores, and repeating this process for 40 transfers. For a filamentous fungus, such as *N. crassa*, that lacks a defined germ line since nuclei can move within the mycelium and all parts of the mycelium can produce asexual spores, the number of MA transfers does not correspond to the number of generations in a biologically meaningful way. Therefore, we express mutation rates per mitosis instead of generation. We used our previous estimate of 25 mitoses per transfer Villalba de la Peña et al. (2023), which corresponds to 1015 mitoses per line for the entire MA experiment, to calculate mutation rates (Villalba de la Peña et al., 2023).

To initially assess DNA methylation divergence, we sampled ten MA lines randomly from two independent MA pedigrees after MA transfers five, twenty, and forty. Strains were grown overnight in liquid culture, DNA was extracted, and bisulfite sequencing was performed from frozen DNA stocks; these protocols are detailed below. We used one replicate per time point for all lines, except for line 10 for which we had two biological replicates from each time point; three biological replicates were performed for both ancestors, for a total of 69 bisulfite sequencing samples. For a more detailed assessment of epimutation rates, we sampled three more MA lines at transfers one, five, seven, eight, ten, and fifteen from both pedigrees for Nanopore sequencing, with one replicate performed for each line, time point, and ancestor, for a total of 38 Nanopore sequencing samples. From four of the Nanopore-seq MA lines, we took samples for H3K9me3 ChIP-seq after transfers five, eight, ten, twenty, and forty; four ChIP-seq replicates were performed for the *mat A* ancestor while two biological replicates were performed for the *mat a* ancestor for a total of 25 ChIP-seq samples.

### DNA extraction and library preparation

Strains to be sequenced were inoculated into a liquid VM culture (Metzenberg, 2003) with 1.5% sucrose and were grown overnight at 25 *^◦^*C. Mycelium was harvested, and high quality DNA was extracted using a phenol-chloroform extraction as in Villalba de la Peña et al. (2023). DNA samples were spiked with 1 ng of unmethylated *λ*-phage DNA (Ambion, USA) to monitor bisulfite conversion efficiency.

For whole genome bisulfite sequencing, DNA samples were sent to Novogene (Cambridge, UK). For library preparation, DNA was sheared into 200–400 bp fragments, and single stranded fragments were treated with bisulfite using the Accel-NGS Methyl-Seq DNA Library Kit for Illumina (Swift Biosciences, USA), per the manufacturer’s protocol. Methylation sequencing adapters were ligated, and Illumina sequencing libraries were size selected within a 200 to 800 bp range and PCR amplification was performed per the manufacturer’s protocol. Methylation-sequencing libraries were sequenced on Illumina machines to generate 150 bp paired-end reads.

Library preparation for Nanopore sequencing was performed using the Native Barcoding Kit 24 V14 (Oxford Nanopore, UK), and DNA was sequenced using MinION flow cells (Oxford Nanopore), per the manufacturer’s instructions.

### Chromatin immunoprecipitation and library construction

H3K9me3-specific ChIP-seq was performed as previously described (Jamieson et al., 2013; Klocko et al., 2019; Scadden et al., 2023), see the supplementary methods for details. ChIP-seq library barcoding was performed using the NEBNext Ultra II kit for Illumina (New England Biolabs, USA) per the manufacturer’s protocols except that eight PCR cycles were used for the library amplification to minimize AT-rich DNA depletion (Ji et al., 2014). ChIP-seq libraries were sequenced on an Illumina NovaSeq 6000 (Genomics and Cell Characterization Core Facility, University of Oregon).

### Bioinformatic analyses

#### Read mapping and cytosine methylation calling from bisulfite sequencing data

An overview of the workflow is shown in Figure S27. The bisulfite conversion protocol used in library preparation adds low complexity nucleotide tails to 3’ ends of each fragment, which appear in the beginning of read 2. These tails were trimmed by fastp v. 0.23.2 (Chen, 2023) with the parameter “trim-front 10”. Trimmed reads were mapped to an *in silico* bisulfite converted reference genome using Bismarck v. 0.22.3 (Krueger and Andrews, 2011) with the dovetail option. Methylation level was called for each cytosine using Bismarck methylation extractor, and the METHimpute R package v. 1.8.0 was used to generate a methylome for each sample. METHimpute, which uses a Hidden Markov Model to classify each cytosine in the genome to either methylated or unmethylated state (Taudt et al., 2018), was then used for calling cytosine methylation status.

Table S1 shows reads, depth, coverage, and bisulfite conversion efficiency estimated from *λ*-phage DNA for each sample.

#### Basecalling, read mapping, and methylation calling from Nanopore sequencing data

Nanopore data was basecalled with the Dorado (v. 0.3.3) basecaller using the super accurate model (dna_r10.4.1_e8.2_400bps_sup@v4.2.0) for identifying each base; this protocol concurrently called modified cytosines. Dorado was also used for sample demultiplexing and mapping the reads to the reference genome, using the minimap2 algorithm (Li, 2018). Methylated sites were called using the modkit (Oxford Nanopore) program, and modkit output was processed with METHimpute to generate a methylome for each sample. Table S2 shows number of reads, their lengths, and depth for each sample. For calling genetic mutations from the Nanopore data, see supplementary methods and Figure S28.

#### Bioinformatic analyses of ChIP-seq datasets

ChIP-Seq reads were aligned to the *N. crassa* reference genome version 12 using BWA or bowtie2 (Li and Durbin, 2009; Langmead and Salzberg, 2012). Output sam files were converted to bam files, sorted, and indexed with samtools (Danecek et al., 2021), and made into Reads per Kilobase per Million Reads (RPKM) 500 or 5000 basepair bin bedgraph files with deeptools (Ramírez et al., 2016) for display with Integrative Genomics Viewer (Thorvaldsdóttir et al., 2013). Enriched ChIP-seq sites were identified using MACS2 (Zhang et al., 2008). First, duplicate reads were removed, as they are likely to result from technical bias. Next, we estimated the fragment length for each sample individually, which varied slightly between samples (ranging from 240 to 298 bp). Fragment length is defined as the distance between the modes of read density peaks on the positive and negative strands. MACS2 shifts each read by half the estimated fragment length and creates bins along the genome, with a bin size twice the fragment length, to scan for peaks. Each sample was also compared with the control to estimate local and global background noise. The peaks were called using the `-broad` option.

To validate the peaks identified by MACS2, we used ChromHMM (Ernst and Kellis, 2017), which infers the chromatin state signatures using a hidden multivariate Markov model. ChromHMM was run with a bin size of 200 bp and the default parameters for all other settings.

We also investigated whether the H3K9me regions diverged over time. To identify unique peaks among samples, a list of nonredundant peaks was initially generated, and overlapping ranges were calculated to construct a presence/absence matrix with the bioconductor GenomicRanges (Lawrence et al., 2013) package.

We used ChromTime software (Fiziev and Ernst, 2018) to investigate whether the H3K9me regions expanded or contracted over transfers. Most of the changes observed occurred in the centromere. However, these expansions and contractions appear to be artifacts of sequencing and peak calling (see Figure S25 and S26 for examples).

#### Chromatin modifications

The *N. crassa* genomic loci enriched with H3K9me3, H3K27me3, and the centromeric CenH3 histone variant were determined from Jamieson et al. (2013) and Smith et al. (2011) as in Villalba de la Peña et al. (2023).

### Statistical analysis

#### Spontaneous DNA methylation changes

DMRs were estimated from METHimpute output by using jDMR (Hazarika et al., 2021), which uses a Hidden Markov Model to segment the genome into 100 bp regions, and looks for nearby cytosines that change their methylation status together to infer the methylation status of each segment. Segments that differed in their methylation status among the samples were classified as DMRs. The minimum number of methylated cytosines present in a segment was set to five cytosines, with each cytosine requiring a minimum read depth of five reads, for a segment to be classified as a DMR.

Methylation divergence among the samples for the bisulfite sequencing data and for the nanopore data was calculated using AlphaBeta (Shahryary et al., 2020). Divergence was calculated from METHimpute output for both single sites and DMRs in CG, CHG, and CHH contexts. We used the number of mitoses as the unit of divergence in the MA pedigree. Because *N. crassa* is haploid, epimutation rate estimates were computed using the AlphaBeta neutral SOMA model, which was modified to account for haploidy (see supplementary methods for details). The neutral model was tested against a null model of no accumulation using a modified model comparison function of AlphaBeta. We used 100 starts for the numerical estimation procedure. Intervals for the estimates were obtained by bootstrapping using 1000 bootstrap replicates.

#### Analysis of H3K9me3 divergence

For the analysis of divergence (e.g., the presence and absence of peaks), we calculated divergence as

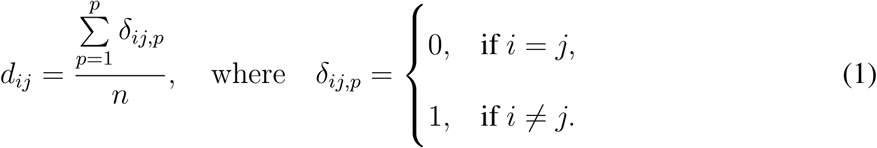

Here *d_ij_*is the divergence between samples *i* and *j*, *δ_ij,p_* is a function that takes a value of 0 if both samples are in the same state, and a value of 1 if they are in a different state for the *p*th peak, and *n* is the number of peaks.

For the analysis of H3K9me3 read counts, we divided the genome into 5kb bins and the centromeric regions into 500bp bins using bedtools and intersected this with ChIP-seq bam files for H3K9me3 to give levels of H3K9me3 for each window across the genome. We then normalized the read counts, and calculated divergence between each sample pair within a pedigree as

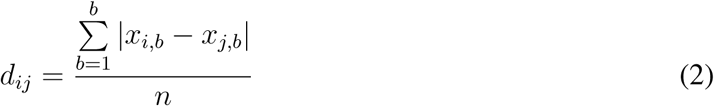

where *d_ij_* is the divergence between samples *i* and *j*, *x_i,b_* is the read count of *ith* sample of the *b*th bin, and *n* is the number of bins. See the supplementary methods for details.

#### Data access

The genome sequencing data generated in this study has been submitted to the European Nucleotide Archive database (https://www.ebi.ac.uk/ena/) under project accession number PRJEB108830. All H3K9me3 ChIP-seq data have been submitted to the NCBI Gene Expression Omnibus database (https://www.ncbi.nlm.nih.gov/geo/) under accession number GSE313506. Other data and scripts are available at https://github.com/ikron/epimutation and as supplemental material.

## Acknowledgments

This study was supported by a grant from the Research Council of Finland (no. 321584) to IK, and a National Institutes of Health (NIH) R15 AREA award (R15140396) to ADK. We thank the Finnish CSC-IT Center for Science Ltd. for providing computational resources.

## Author contributions

I.K., M.V., F.J., M.C.-T., A.D.K. conceived the study. M.V., C.H.-C., I.K. performed experiments. M.V, I.K., C.H.-C., T.R.H, C.A.V., A.D.K., P.S., F.J. analyzed the data. I.K. and M.V. wrote the manuscript. All authors edited the final manuscript.

