## Supplementary Information for "Centromeres are hotspots of cytosine methylation epimutations in a filamentous fungus"

### 1 Supplementary Information

#### 1.1 Supplementary methods

##### 1.1.1 Chromatin immunoprecipitation

Strains were grown in VM media overnight at 32 °C. Mycelial pellets were harvested with a Büchner filter, crosslinked with 1% formaldehyde (Sigma Aldrich) in PBS for 15 minutes at room temperature, washed with PBS, and sonicated in ChIP Lysis Buffer (50 mM HEPES, pH 7.5; 140 mM NaCl; 1 mM EDTA; 1% Triton X-100) and Halt Protease and Phosphatase Inhibitor Cocktail (Thermo Fisher Scientific, # 78441), for concurrent cell lysis and chromatin shearing in a Bioruptor Pico (Diagenode), with a 15 minute cycle of 30 seconds sonicating 30 seconds off. The $\alpha$-H3K9me3 antibody (ActiveMotif cat# 39161) was added to cleared lysate and incubated overnight at 4 °C. Protein A/G magnetic agarose beads (ThermoFisher, cat# PI78609), washed with ChIP Lysis Buffer, were added to the lysate and incubated for two hours at 4 °C. Beads were subsequently washed twice with ChIP Lysis Buffer, once with ChIP Lysis Buffer + 0.5 M NaCl, once with LiCl wash buffer (10 mM Tris-HCl pH 8.0; 250 mM LiCl; 5% IGEPAL CA-630; 1 mM EDTA), and DNA was eluted with 125 µl of TES buffer (50 mM Tris-HCl pH 8.0, 10 mM EDTA, 1% SDS). ChIP-DNA was treated with 100 µg/ml RNaseA, 500 µg/ml Proteinase K, extracted once with phenol/chloroform/Isoamyl alcohol (25:24:1), once with chloroform, ethanol-precipitated with 160 µg/ml glycogen added for DNA recovery, and the ChIP-DNA pellet was resuspended in 30 µl TE. The final ChIP-DNA sample was quantified with a Qubit 3.0, using the HS method.

##### 1.1.2 Analysis of H3K9me3 divergence

For the analysis of H3K9me3 divergence using read counts, we first removed all bins that were not in the seven chromosomes of assembly NC12. Then we normalized the read counts following Love et al. (2014). First we calculated geometric mean for each bin, $g_b$ across all samples as

$$g_b = \exp\left(\frac{\sum_{i=1}^{i} \ln x_{b,i}}{n}\right)$$

where $x_{b,i}$ is the read count of $b$th bin for sample $i$, and $n$ is the number of samples. Then a size factor was calculated for each sample by first dividing the count of each bin and each sample by $g_b$ of the $b$th bin, and then taking the median from all ratios for each sample: $s_i = \text{med}\,(x_{b,i}/g_b)$. Then normalized count, $x_{n,i}$ is obtained by dividing each count by the $s_i$ of that sample, as $x_{n,i} = x_i/s_i$. Then the normalized counts were used in the divergence analysis.

##### 1.1.3 Haploid model for estimating epimutation rates

The original model of AlphaBeta assumes diploid data (Shahryary et al., 2020). Since *N. crassa* is haploid we modified the AlphaBeta somatic model to account for haploidy. Calculating divergence remains the same as in the diploid model. Divergence gets a value of 1 if the cytosines or regions being compared differ in their methylation status, and 0 otherwise. What changes from the diploid model is the transition matrix, $\mathbf{G}$, which in the case of haploid data simplifies to

$$\mathbf{G} = \begin{array}{c} \\ c^u, t \\ c^m, t \end{array} \begin{array}{c} \begin{array}{cc} c^u, t+1 & c^m, t+1 \end{array} \\ \begin{pmatrix} 1-\alpha & \alpha \\ \beta & 1-\beta \end{pmatrix} \end{array}$$

where $c^u$ and $c^m$ are unmethylated and methylated states respectively, $\alpha$ is the rate of methylation, and $\beta$ is the rate of methylation loss. The proportions of methylated and unmethylated cytosines when the system is at dynamic equilibrium, $\pi = (\pi_u, \pi_m)$, can be found using Markov chain theory $\pi = \pi\mathbf{G}$. This implies a system of linear equations

$$(1-\alpha)\pi_u + \beta\pi_m = \pi_u$$

$$\alpha\pi_u + (1-\beta)\pi_m = \pi_m$$

$$\pi_u + \pi_m = 1$$

that can be solved by substituting $\pi_m = 1 - \pi_u$ and solving for $\pi_m$ and $\pi_u$:

$$\pi = \left( \frac{\alpha}{\alpha + \beta}, \frac{\beta}{\alpha + \beta} \right).$$

The equilibrium proportions of methylated and unmethylated cytosines depend only on rates of methylation gain and loss.

The haploid model also has one parameter less than the diploid model, as there is no proportion of epiheterozygotes in the model. Estimation of the rate parameters was done as in Shahryary et al. (2020), with numerical estimation using the Nelder-Mead algorithm.

##### 1.1.4 Calling genetic mutations from the Nanopore data

Four our previous work, we know the spontaneous genetic mutations that had happened in the final MA transfer. These mutations were called using Illumina sequencing, and we have validated genotyping pipeline using different methods, including validating a set of the mutations by Sanger sequencing (Villalba de la Peña et al., 2023). To call these mutations from the intermediate MA transfers from the Nanopore-seq dataset we simply examined the read alignment of each sample in IGV. Our observation is that when Nanopore basecalling has been performed using the super accurate model in Dorado, real SNPs can be identified from the alignments unambiguously. Indeed, when we compared the Illumina alignment from Villalba de la Peña et al. (2023) with the Nanopore alignments at positions where the known mutations occur, we could identify the transfers where this mutation appeared (Figure S28).

#### 1.2 Supplementary results

##### 1.2.1 Analysis of divergence outliers in WGBS data

We initially observed that the variance in divergence among the WGBS samples increased substantially after approximately 500 mitoses, particularly for single cytosines (Figure 2). We examined the possibility that this phenomenon was caused by one or few samples that were outliers, due to a technical issue in the library preparation for example, and all pairwise comparisons to such outlier samples were biased as a result. We first looked at the data for single cytosines in CG context: we chose an arbitrary threshold of divergence of 0.00175 as bulk of the data seemed lie below this threshold, and asked were there particular samples that were associated with pairwise comparisons that exceeded this threshold (Figure S3). We identified four samples out of 69 that were associated with pairwise comparisons with extreme values: Line 13 transfer 20, Line 6 transfer 40, Line 35 transfer 20, and Line 36 transfer 40 (Figure S3). Removing these samples from the data removed nearly all pairwise comparisons with extreme values. However, increase in variance after 500 mitoses could still be observed even after these outlier samples were removed (Figure S3).

We then fitted an AlphaBeta model of epimutation accumulation to the data where we had excluded the four outlier samples described above (Figure S4). Removing these outlier samples does reduce the variance compared to the full dataset, particularly for DMR divergence (Figure 2). However, in all cases the model still does not fully capture the saturation effect in the data (Figure S4). A model of neutral accumulation of epimutations is preferred over null model of no accumulation for single cytosines: $p < 2.2 \times 10^{-16}$ in CG, CHG, and CHH contexts. The results were similar for DMRs: the model of neutral epimutation accumulation was preferred over the null model: $p < 2.2 \times 10^{-16}$ in all sequence contexts.

###### 1.2.2 Robustness of results to DMR specification

In the original analysis we performed DMR segmentation by binning the genome into 100 bp windows and requiring a minimum of 5 cytosines to change their methylation status together. Of course choosing the window size and number of cytosines, is somewhat of an arbitrary choice. Therefore, we checked whether the results were robust to different choices of parameter values for DMR segmentation. In addition, we performed segmentation for 200 bp windows, requiring a minimum of 10 cytosines to change their methylation status, and for 300 bp windows with 12 cytosines. We observed that these choices did not substantially change the biological results. We still observed a clear signal of DMR enrichment in the centromeric regions in the WBGS dataset

(Figure S5A). The only difference was that there were less longer DMRs in regions of interspersed heterochromatin, so regions marked by H3K9me3 that were not in the centromere (Figure S5A).

Furthermore, we tested whether the methylation divergence observed for DMRs in the MA lines in centromeric regions in the Nanopore dataset was robust to DMR specification, and we observed that different window sizes had no effect on divergence (Figure S5B). This is perhaps unsurprising as the divergence is observed for single cytosines as well (Figure 4).

We chose to present the analyses mainly for 100 bp binning. The argument that favors this choice is that it allows the most fine grained analysis of methylation changes which can allow to detect small scale shifts in methylation patterns. One could argue that since we know that DNA methylation in *N. crassa* requires H3K9me3, thus the length of the nucleosome, which is approximately 200 bp, or longer could also be a natural choice for window size. However, by setting the window smaller than this, we can potentially capture events where the position of a nucleosome changes slightly. Nevertheless, our main results regarding DMRs, that they are enriched in the centromeric regions, and that the MA lines diverge from each other in DMR patterns in the centromeric regions, are robust to parameter choices in the DMR segmentation.

##### 1.2.3 Changes in centromeric methylation proportion in the MA lines

We examined whether the observed divergence in methylation patterns in centromeric H3K9me3 could be caused by just the centromeres losing methylation over time. First we examined how many cytosines in the centromeric regions were called as methylated in the MA lines, and plotted proportion of methylated cytosines in the centromere for each MA line over the MA transfer. We observed that in the WGBS dataset there was a tendency for methylation to generally decrease over the experiment, although there were a few MA lines where methylation levels remained constant or slightly increased (Figure S11A). Methylation proportion followed the same patterns regardless of sequence context (Figure S11). However, in the Nanopore dataset, where our resolution over the transfer was higher, we observed again that overall methylation proportion in the centromere decreased for four out of the six lines and increased for the remaining two lines (Figure S11B).

For the Nanopore dataset, where we had dense sampling of the MA transfers, we examined consecutive time points and determined how many gain and loss of methylation events happened for each transfer interval. Detailed analysis of specific intervals per line is shown in Figure 5, and in Figure S15. Figure S11C shows a summary of methylation gains and losses across all transfer intervals and MA lines. There are a substantial number of gain and loss events for both DMRs and single cytosines. For single cytosines, methylation losses were more common, with 382 876 losses and 375 567 gains of single cytosine methylation. For DMRs we observed 45 124 gains of methylation and 42 085 losses. This shows that while methylation loss is more common, and accordingly our estimate of loss rate is higher (Table 1), methylation patterns in the centromeres do not diverge only because of methylation is gradually lost, but methylation gains happen constantly and shape the observed divergence. Methylation in the centromeres is eventually expected to reach dynamic equilibrium.

##### 1.2.4 Methylation divergence and nucleosome position drift

One hypothesis that could explain divergence in methylation patterns without invoking epimutations, is drift in nucleosome positions. In *N. crassa* H3K9me3 is required for DNA methylation, thus if it is so that that each cytosine that is in contact with the nucleosome is going to be methylated, and those cytosines that are not in contact with the nucleosome cannot be methylated, then it could be possible that DNA methylation patterns could change if nucleosomes are placed at slightly different positions after each round of DNA replication. This kind of nucleosome position drift would make the MA lines diverge from each other.

However, if DNA methylation divergence follows this model, this should be observed as a specific pattern of DNA methylation. It has been estimated that 146–147 bp DNA is wrapped around the histone octamer, and the nucleosomes are connected by 20–60 bp of linker DNA. Thus, the length of one nucleosome is about 170–200 bp, perhaps closer to the shorter range in the compacted centromeric regions. The implication is that if DNA methylation divergence is caused by nucleosome position drift, then DNA methylation losses and gains should always happen at very short scales. We always observed a very strong signal of H3K9me3 enrichment in the centromeric regions, with no differences among the MA lines. Therefore there cannot be large scale differences in H3K9me3, and that only leaves the possibility of short scale changes in nucleosome positions that are below the Chip-seq read resolution.

To evaluate this hypothesis we examined DNA methylation patterns in the MA lines. We took the inferred DMR positions, and then looked at all methylation, including invariable sites, in these regions for the different time points from the Nanopore dataset, where we had denser sampling. We observed different kinds of methylation patterns, most notably some methylation changes that spanned much longer stretches of DNA than nucleosome length. For instance Figure S12A shows a locus that was unmethylated in the MA ancestor, but at transfer 5 MA line 5 had gained some methylation, which increased further in in transfer 7 and 8. Yet, by transfer 15 this methylation was subsequently lost again. The methylated regions spans approximately 1 kb in length, and there was no methylation present in immediate upstream or downstream regions. This does not seem compatible with a model where the position of nucleosome slightly shifts to produce this pattern. Similarly, Figure S12B shows an example in MA line 23, where an approximately 2 kb regions shows variable methylation patterns up to transfer 5, then majority of methylation is lost in transfer 7 and 8. However, by transfer 15 this region is remethylated. This again seems incompatible with short range movement of nucleosomes.

There are instances of methylation changes which could be compatible with nucleosome movement. For instance, Figure S12C and S12D show examples where DNA methylation seem to be spreading into an upstream unmethylated region, which is flanked by a region of constant invariable methylation. However, it is not clear if these examples really represent nucleosome movement, since we don't observe a corresponding loss of methylation downstream of the spreading methylation (Figure S12C and S12D). Thus, it seems that even in these cases a simple model of nucleosome position drift does not explain these patterns.

In total we observed tens of thousands of gain and loss of DMR events (Figure S11) in the MA pedigrees. We of course cannot exclude that some of these events could be due to local changes in nucleosome position. However, taken together, the patterns of methylation divergence we observe in the centromeric regions represent gains and losses of methylation that are not due to solely just methylation changes brought about by local changes in nucleosome positions, but other mechanisms must be involved. Therefore, the DNA methylation divergence we observe does reflect genuine epimutations that happen and are transmitted in the centromeric regions.

##### 1.2.5 DMRs in different classes of transposable elements

We investigated whether different types of TEs in centromeric and non-centromeric regions could in principle explain why centromeric regions have so much more DMRs. It could be that TE composition in centromeric and non-centromeric region is different and the TE classes present in centromeric regions are especially prone for DNA methylation changes. To examine this possibility, we first checked which classes of TEs are present in centromeric and non-centromeric regions (Figure S6A). There were some differences in TE composition of centromeric and non-centromeric regions: DNA transposons were more common in non-centromeric regions, even this class is very rare in the *N. crassa* genome. Furthermore, SINEs and unknown elements were also more common in non-centromeric regions, with TEs of unknown class representing the largest difference of 18.1 percentage points between non-centromeric and centromeric regions (Figure S6A). LINE/Tad1 elements, LTRs, Helitrons, and non-LTE elements were common in centromeric regions. However, the difference between centromeric and non-centromeric regions for the second most common class of LTR elements was only 6.6 percentage points, and 11.3 percentage points for LINE/Tad1 elements (Figure S6A).

The second step was to examine whether DMRs were more common in certain TE classes, we fit a model DMR occurrence rates for the different TE classes across the genome. We did observe that DMRs occurred more often in some TE classes relative to random expectation. DNA transposons had lower rates of DMR occurrance than expected based on their abundance, and the same was true for SINE elements (Figure S6B). DMRs that had higher rates of occurrance in LINE/Tad1, Helitrons, and non-LTR class elements, while DMRs occurred at expected levels in LTR elements

(Figure S6B).

To assess whether these differences in TE composition between centromeric and non-centromeric regions and differences in DMR occurrence in different TE classes could explain the differences in amounts of observed DMRs in TEs between centromeric and non-centromeric regions, we used the previous model of DMR rates in different TE classes to predict the amounts of DMRs in centromeric and non-centromeric TEs, given the TE composition of these regions. The model did not predict the observed DMR counts correctly (Figure S6C). This means that much higher amount of DMRs observed in centromeric TEs cannot be explained by TE composition differences alone, and we need to invoke some other centromere specific factor.

###### 1.2.6 Initial DNA methylation divergence in the WGBS data for centromeric regions

While we consider that we get the most precise estimates of epimutation rates from the Nanopore-seq dataset, it is nevertheless interesting to check whether we obtain comparable estimates from the WGBS dataset. Since the saturation effect was not captured well by the epimutation model, we examined only the initial 300 mitoses where divergence in the centromeric regions looks more linear in the WGBS dataset (Figure S7 and S8).

For single sites divergence increases in a nearly linear fashion for the first 300 mitoses (Figure S13). Neutral epimutation accumulation model was preferred over the null model of no accumulation in all sequence contexts: $p = 3.29 \times 10^{-6}$ for CG, $p = 8.93 \times 10^{-11}$ for CHG, and $p = 1.67 \times 10^{-6}$ for CHH context. The epimutation rate estimates we obtained for single sites were roughly comparable to the estimates we obtained from the Nanopore-seq dataset. For CG context we obtained $\alpha = 1.29 \times 10^{-5}$ and $\beta = 6.16 \times 10^{-5}$, so $\alpha$ is consistent with Nanopore dataset and $\beta$ is slightly lower. For CHG context we obtained $\alpha = 1.40 \times 10^{-5}$ and $\beta = 7.79 \times 10^{-5}$, which are both consistent with Nanopore dataset estimates. For CHH context we obtained $\alpha = 1.03 \times 10^{-5}$ and $\beta = 5.85 \times 10^{-5}$, so $\alpha$ is lower than our Nanopore dataset estimate while $\beta$ is consistent. Overall, for single sites the epimutation rate estimates are mostly consistent between the two datasets and differences are within measurement uncertainty (Table 1).

For DMRs we observed a somewhat different picture than for single sites, while divergence for CG context is nearly linear for the first 300 mitoses, for CHG and CHH contexts there seems to be more variance (Figure S13 and S7). Neutral epimutation accumulation model was preferred over the null model of no accumulation in CG context ($p = 1.07 \times 10^{-11}$), in CHG context but not as strongly ($p = 1.17 \times 10^{-3}$), and there was no evidence that accumulation was preferred over the null model in CHH context ($p = 0.19$). The epimutation rate estimates for DMRs were generally not consistent with the Nanopore-seq dataset. For CG context we obtained $\alpha = 6.17 \times 10^{-5}$ which is higher than our Nanopore dataset estimate, and $\beta = 25.60 \times 10^{-5}$ which is also higher. For CHG context we obtained $\alpha = 3.98 \times 10^{-5}$ which is higher than our Nanopore dataset estimate, and $\beta = 1.70 \times 10^{-3}$ which is considerably higher than our Nanopore dataset estimate. For CHH we observed that $\alpha = 4.61 \times 10^{-5}$ which is not too far off from our Nanopore dataset estimate, but $\beta = 1.16 \times 10^{-3}$ which is considerably higher than our Nanopore dataset estimate (Table 1). For CHG and CHH contexts we also observed that the model had numerical convergence issues. There seems to be something in the CHG and CHH contexts in the WGBS DMR dataset that causes strange divergence values for these time points, which in turn leads to problems in epimutation rate estimation.

###### 1.2.7 Spontaneous changes in H3K9me3

When we first analyzed divergence in H3K9me3 read counts we observed that divergence had a very slight negative slope (Figure S24A). If changes in H3K9me3 patterns are transmitted across mitoses the MA lines should diverge from each other and slope of divergence should be positive, and a negative slope does not have a clear biological interpretation. We investigated further what could cause this, and when we plotted divergence that each sample had to other samples within a pedigree we observed that there were two samples that had high divergence to all other samples (Figure S24B). Both of these samples were replicates of the mat A ancestor. Two other replicates of this ancestor did not exhibit the same pattern. This suggested that there could be a technical reason instead, perhaps some bias that occurred during library preparation or sequencing. When we excluded these two samples, slope of the divergence was not different from zero (Figure S24C). We also observed that and when we performed the analysis separately for the two different MA pedigrees, with all samples included, we did not observe significant divergence. For the mat $A$ pedigree: $\beta = -0.096$, $p = 0.058$ and for the mat $a$ pedigree: $\beta = -0.004$, $p = 0.817$. Since we observed that DNA methylation changes were transmitted in the centromeric regions, we analyzed the data also by using only 500 bp bins in the centromeric regions, but we did not observed significant different (Figure S24D) nor was there divergence in regions outside the centromeres (Figure S24E).

#### 1.3 Supplementary Figures

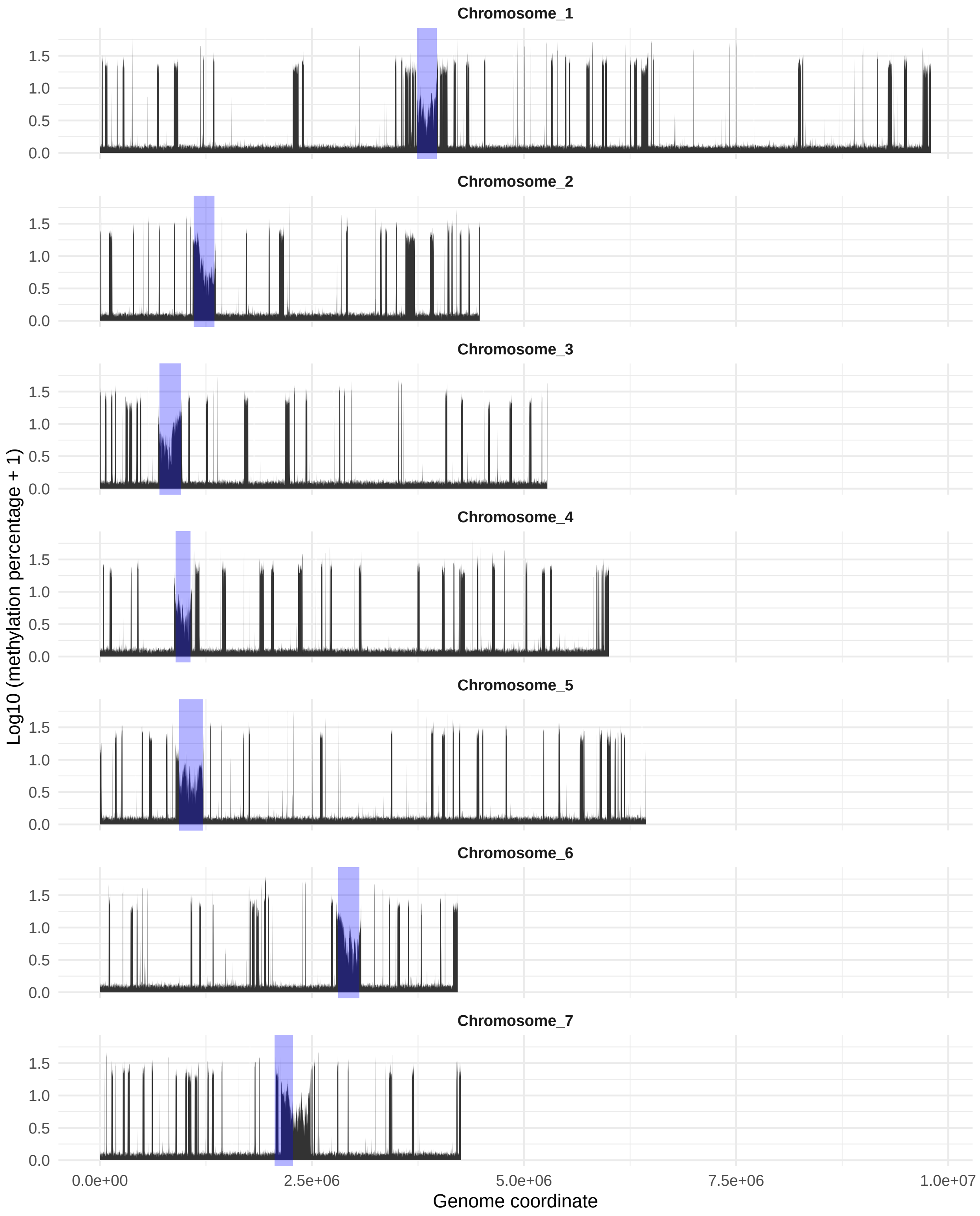

Figure S1: DNA methylation level of the *mat A* ancestor across the seven chromosomes of *N. crassa*. Each chromosome was divided into 1 kb bins, which are represented as genomic positions on the x-axis. The y-axis shows the percentage of DNA methylation (note the $\log_{10}$ scale). The positions of centromeres are shown in blue.

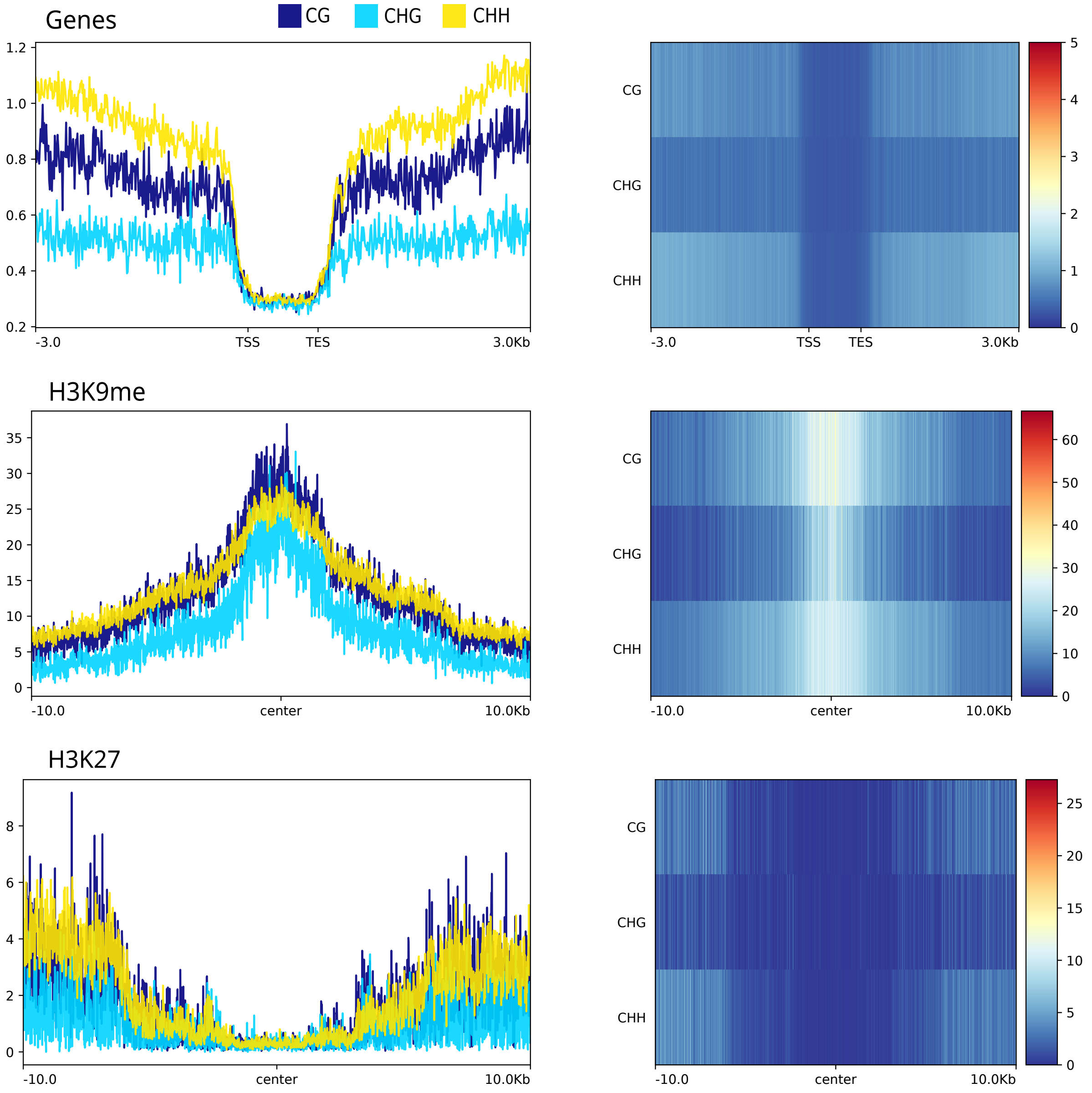

Figure S2: Metaplots and heatmaps showing the percentage of DNA methylation (y-axis) around transcription start sites (TSSs) and transcription end site (TES) (top), within 10 kb of the centers of H3K9me3 regions (middle), and within 10 kb of the centers of H3K27me3 regions (bottom).

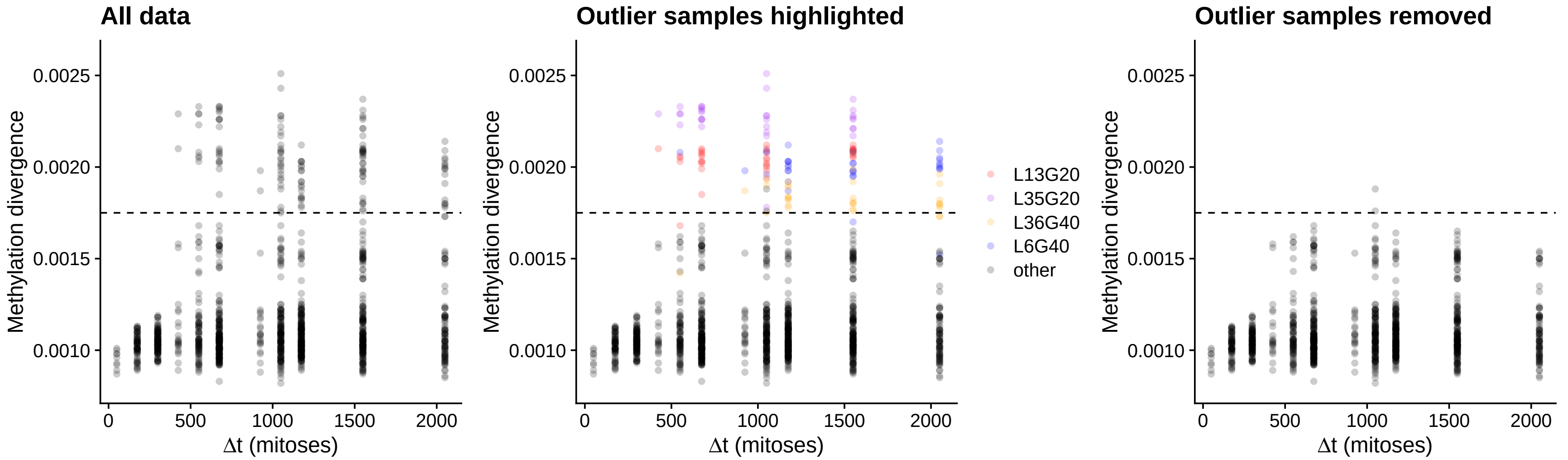

Figure S3: Divergence among the MA lines at single cytosines in CG context for the WGBS dataset. Divergence values above threshold 0.00175, showed by dashed line, were considered outliers. Then samples associated with the outlier pairwise comparisons were identified, and four samples: Line 13 transfer 20, Line 6 transfer 40, Line 35 transfer 20, and Line 36 transfer 40 were observed to be associated with multiple outlier comparisons. Removing these samples from the data removes nearly all extreme divergence values.

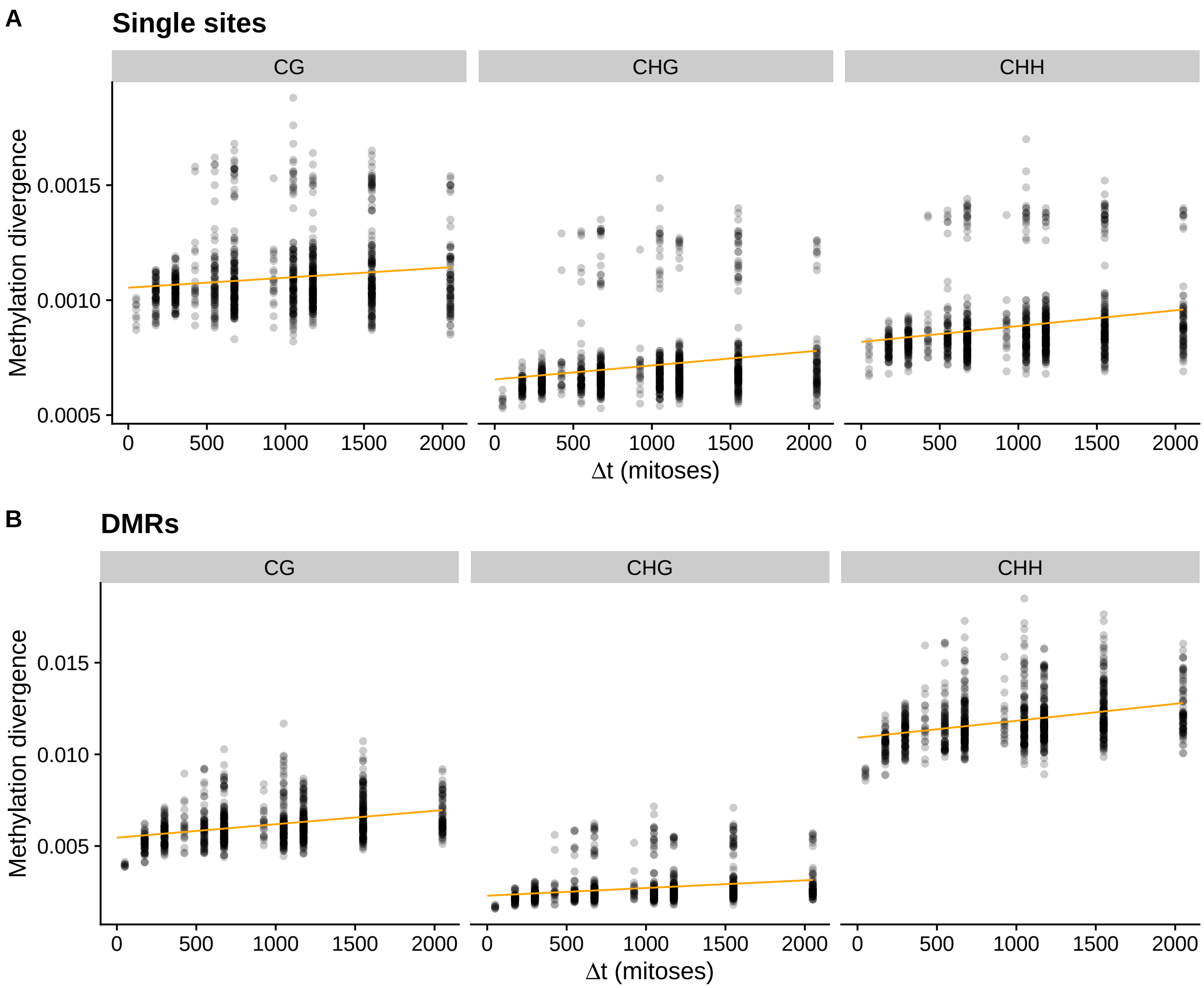

Figure S4: Divergence in cytosine methylation patterns among the MA lines and their ancestors. Outlier samples (Figure S3) have been removed from the data. X-axis shows the number of mitoses separating the two lines that are being compared, and the y-axis shows cytosine methylation divergence. Orange line shows a neutral AlphaBeta model fit. A) The divergence calculated from single cytosines, for CG, CHG, and CHH sequence motifs. B) The divergence calculated for DMRs in different sequence contexts, as in panel A.

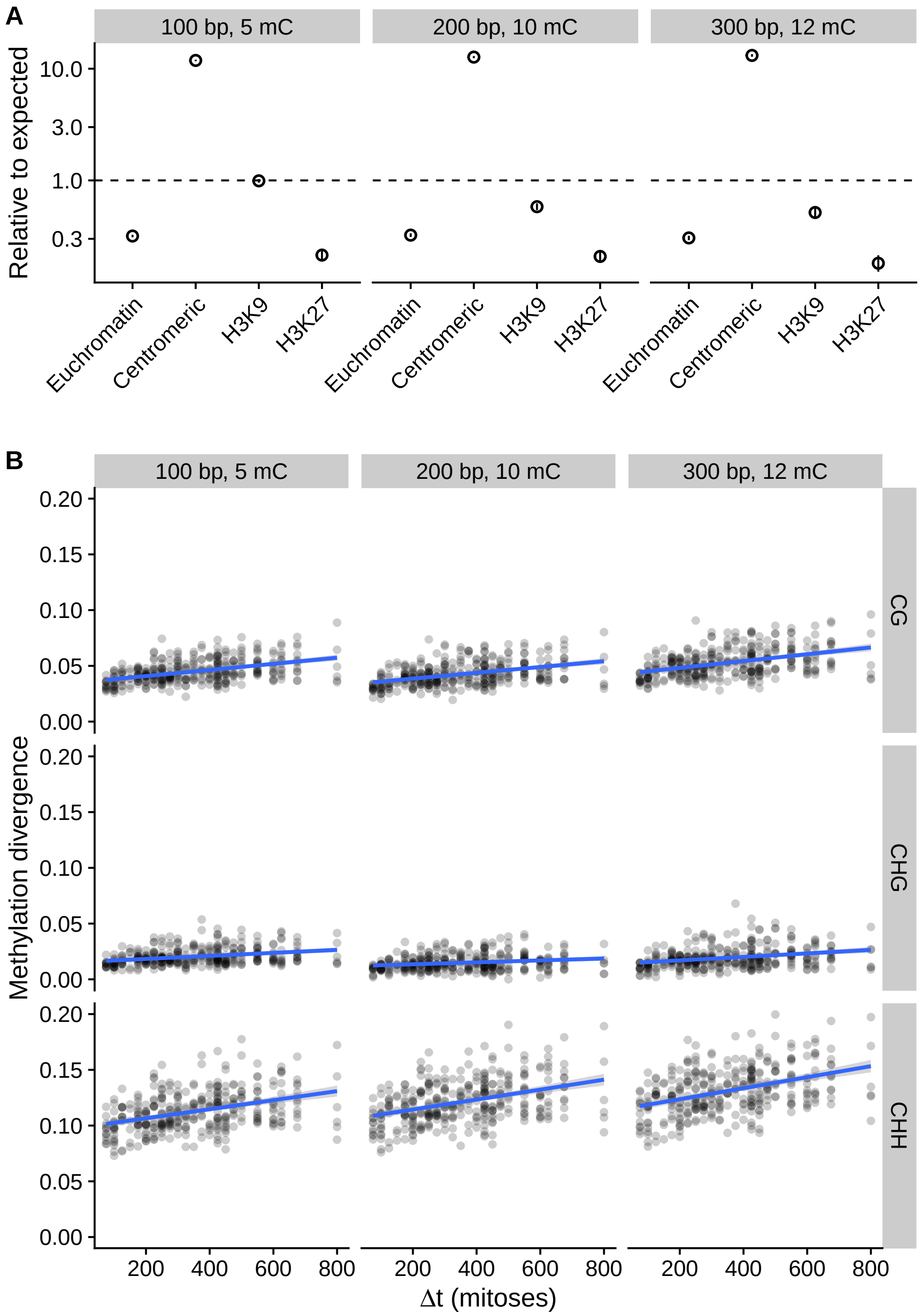

Figure S5: Robustness of results to DMR specification. DMR segmentation was performed for different window sizes and minumum number of required cytosines: 100 bp with 5 cytosines (the original analysis), 200 bp windows with 10 cytosines, and 300 bp windows with 12 cytosines. A) Enrichment of DMRs in the different domains in the WGBS dataset. B) Divergence of MA-lines in centromeric regions in the Nanopore dataset.

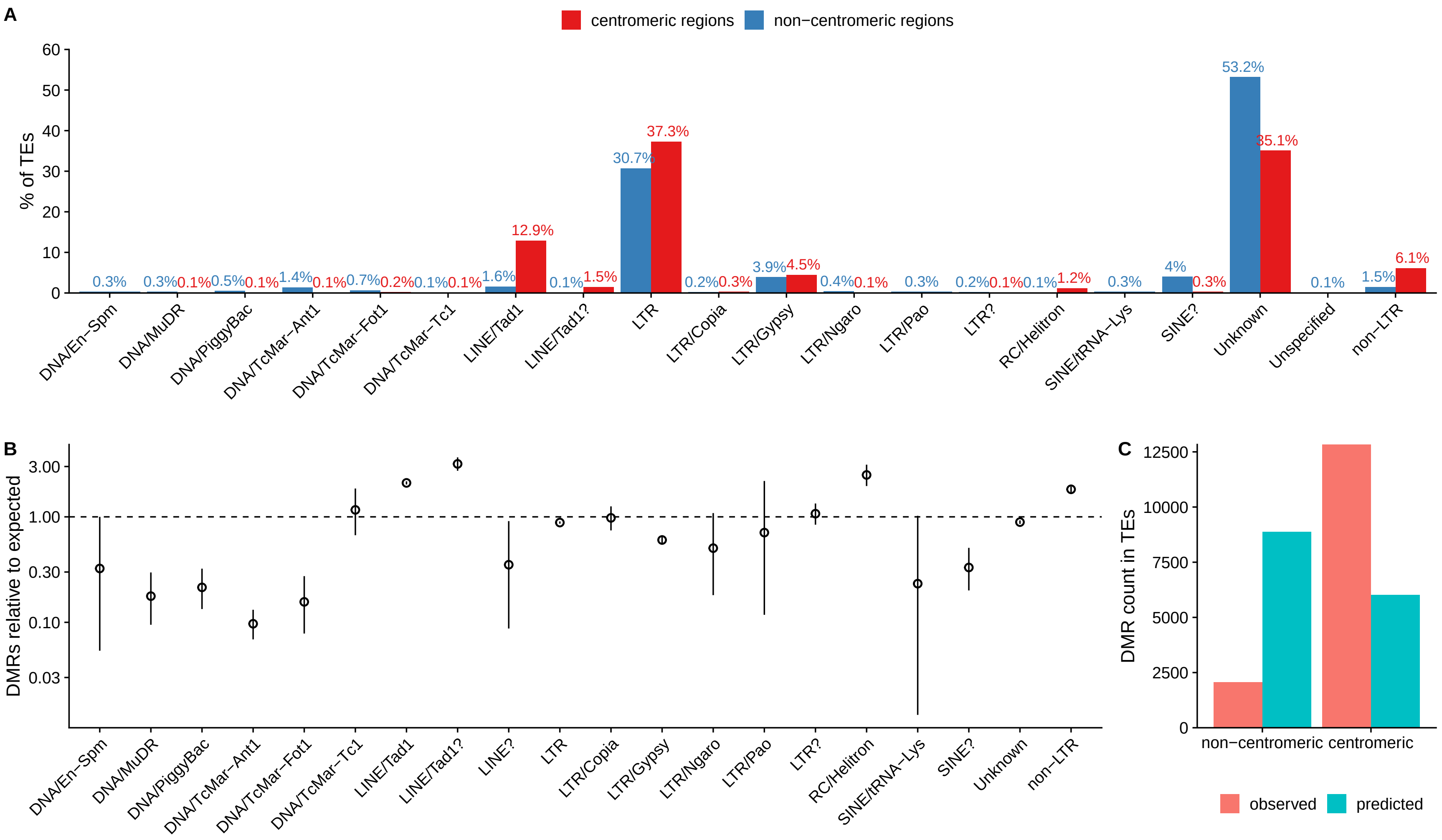

Figure S6: Different TE classes and DMRs. A) Distribution different TE classes among in centromeric and non-centromeric regions. Classes that represented $< 0.1\%$ of TE's were not plotted. B) Rates of DMR occurrence in different TE classes across the whole genome, relative to random expectation. Two exceptionally rare classes of TEs with no DMRs were not plotted as their estimates had large uncertainty. C) Observed and predicted amounts of DMRs, given the distribution of TEs observed in centromeric and and non-centromeric regions (A) and rates od DMRs in these TE classes (B).

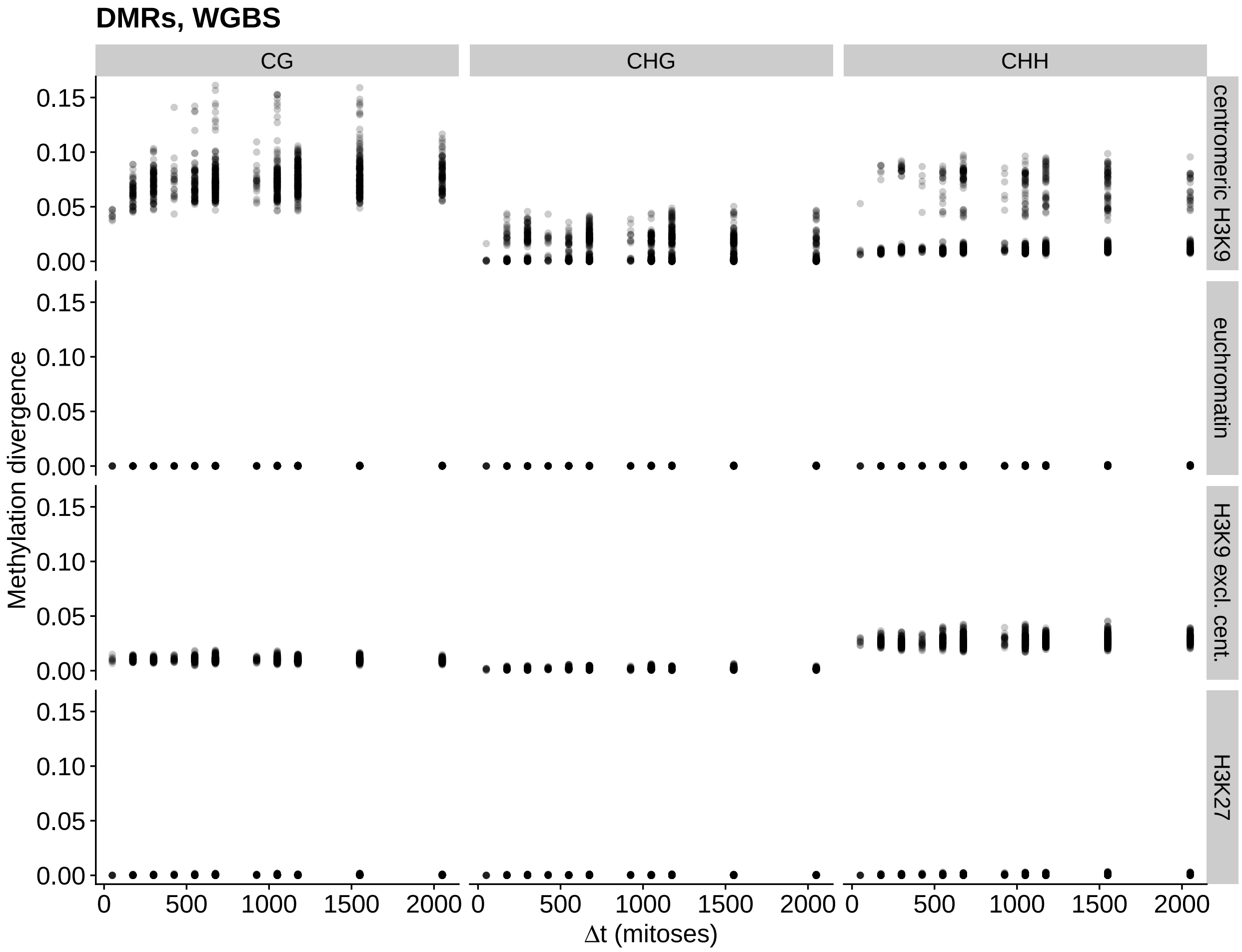

Figure S7: Pairwise divergence in methylation patterns among the MA lines and their ancestors for different regions of the genome. Methylation was detected with WGBS and divergence was calculated for DMRs. X-axis shows the number of mitoses separating the two lines that are being compared, and the y-axis shows methylation divergence.

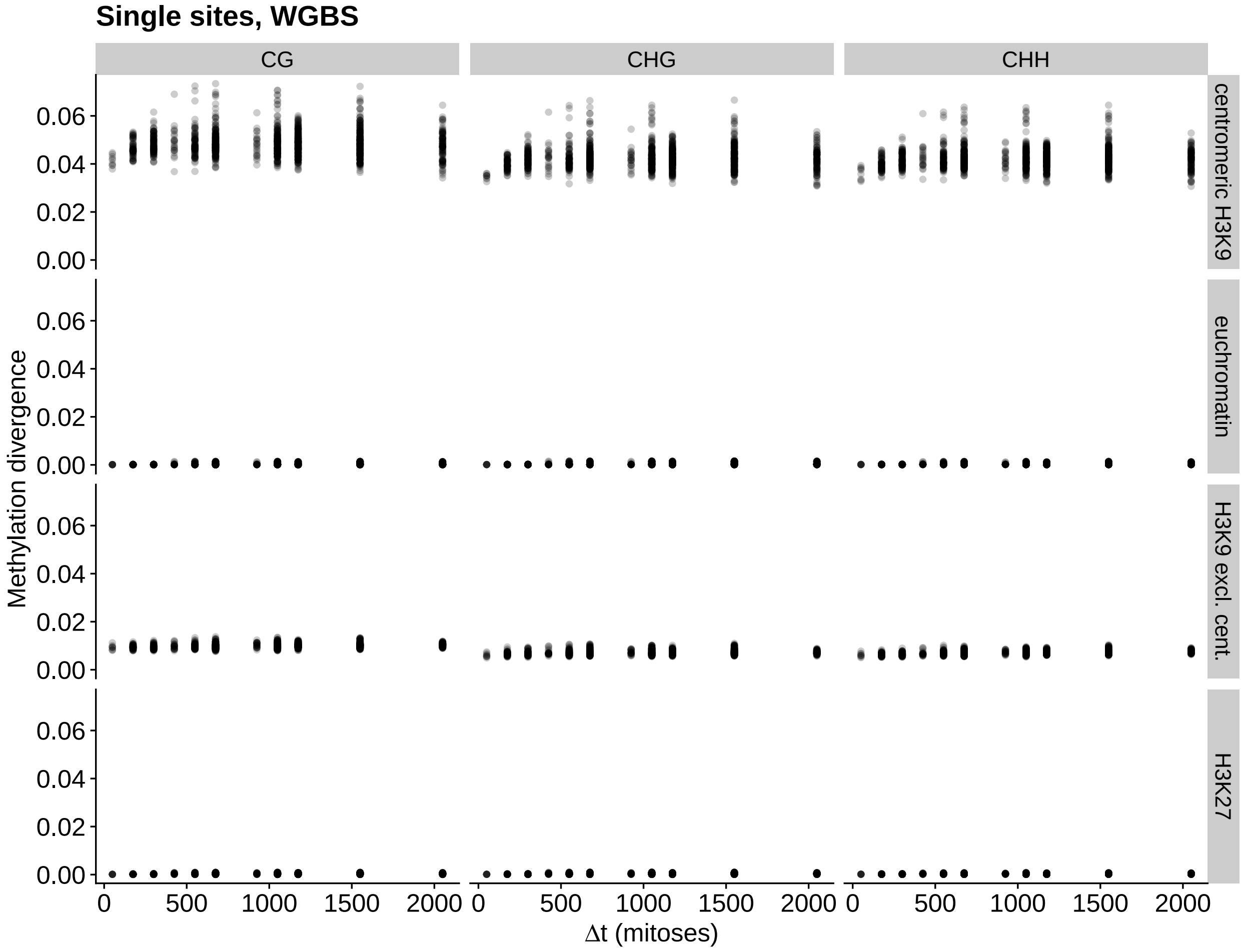

Figure S8: Pairwise divergence in methylation patterns among the MA lines and their ancestors for different regions of the genome. Methylation was detected with WGBS and divergence was calculated for single cytosines. X-axis shows the number of mitoses separating the two lines that are being compared, and the y-axis shows methylation divergence.

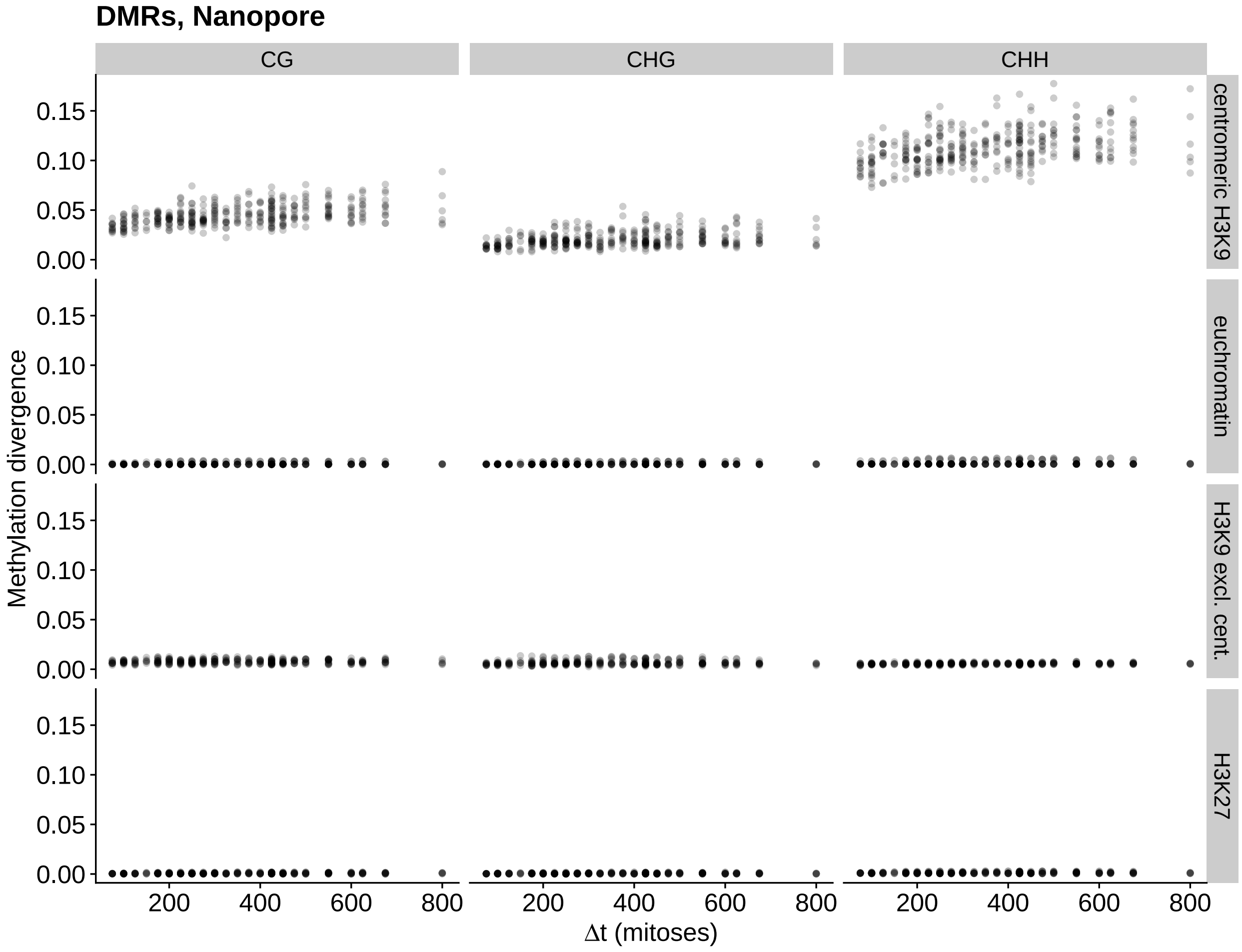

Figure S9: Pairwise divergence in methylation patterns among the MA lines and their ancestors for different regions of the genome. Methylation was detected with Nanopore sequencing and divergence was calculated for DMRs. X-axis shows the number of mitoses separating the two lines that are being compared, and the y-axis shows methylation divergence.

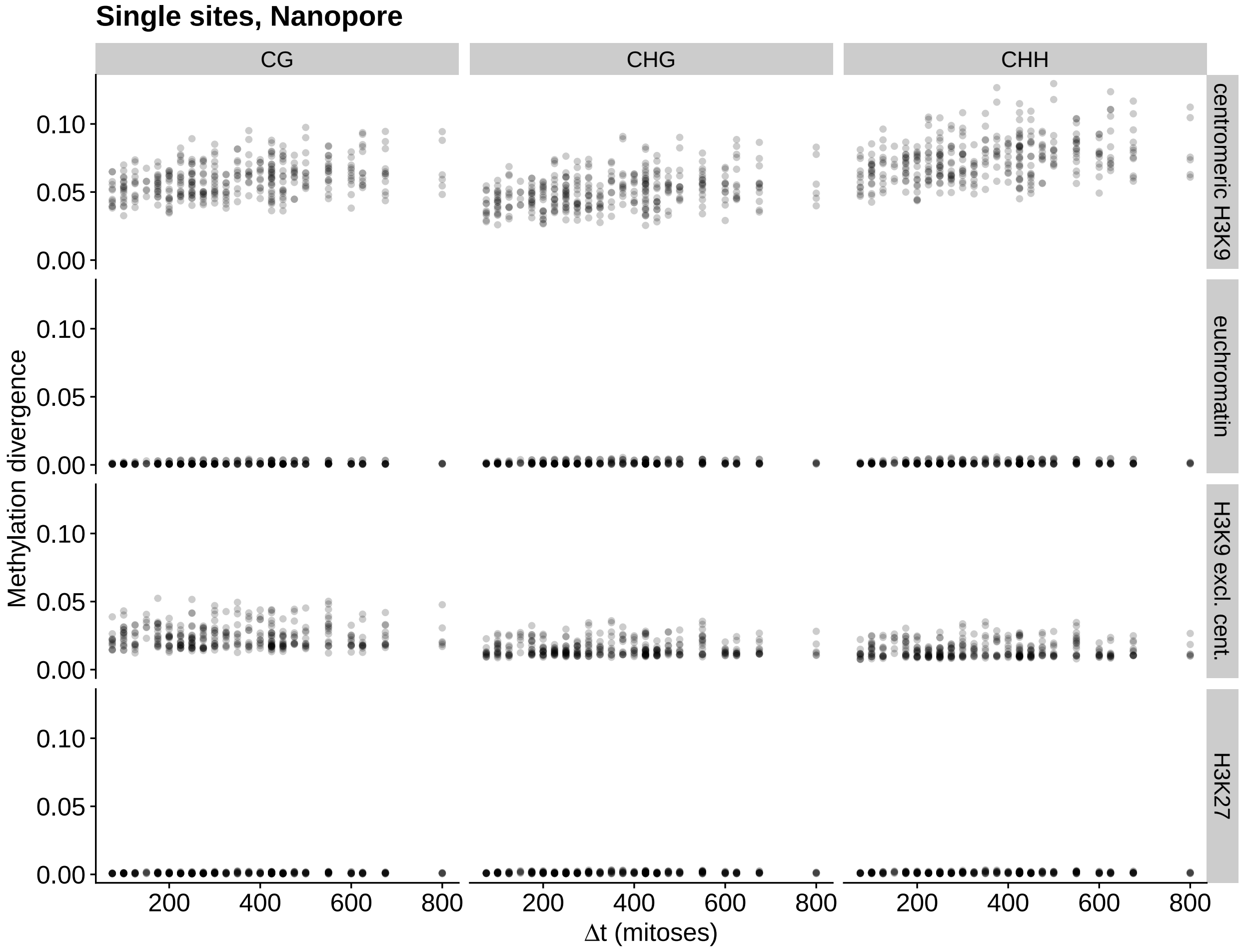

Figure S10: Pairwise divergence in methylation patterns among the MA lines and their ancestors for different regions of the genome. Methylation was detected with Nanopore sequencing and divergence was calculated for single cytosines. X-axis shows the number of mitoses separating the two lines that are being compared, and the y-axis shows methylation divergence.

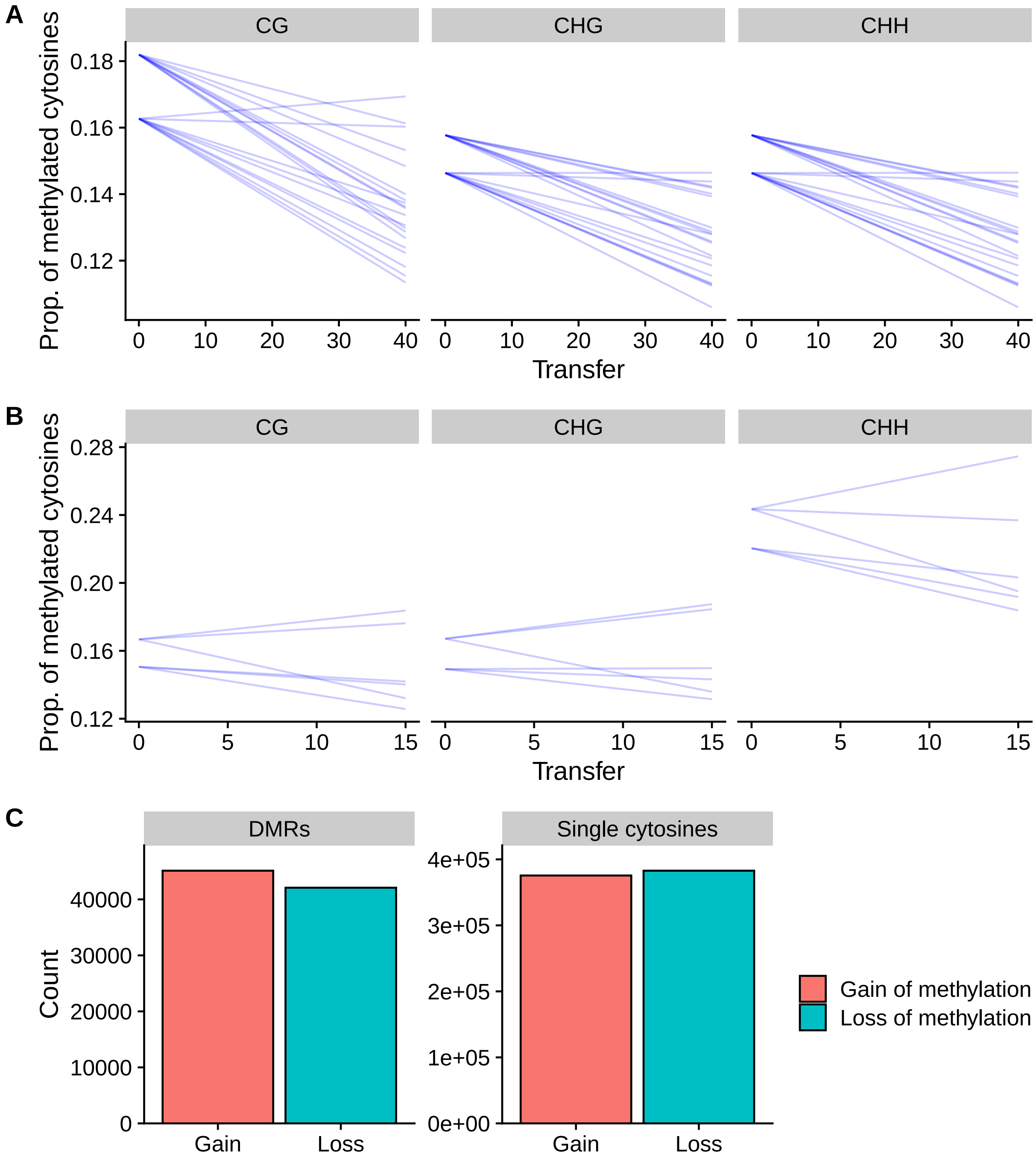

Figure S11: Methylation patterns in centromeric H3K9me3 in the MA lines. A) Proportion methylated cytosines in centromeric H3K9me3 for each MA line, over the transfers for the WGBS dataset. The two starting points, represent the two ancestors of the two MA pedigrees. B) Proportion of methylated cytosines in centromeric H3K9me3 for the MA lines in the Nanopore dataset. C) Total number of gain and loss of methylation events that happened between the different transfer intervals for the MA lines in the Nanopore dataset.

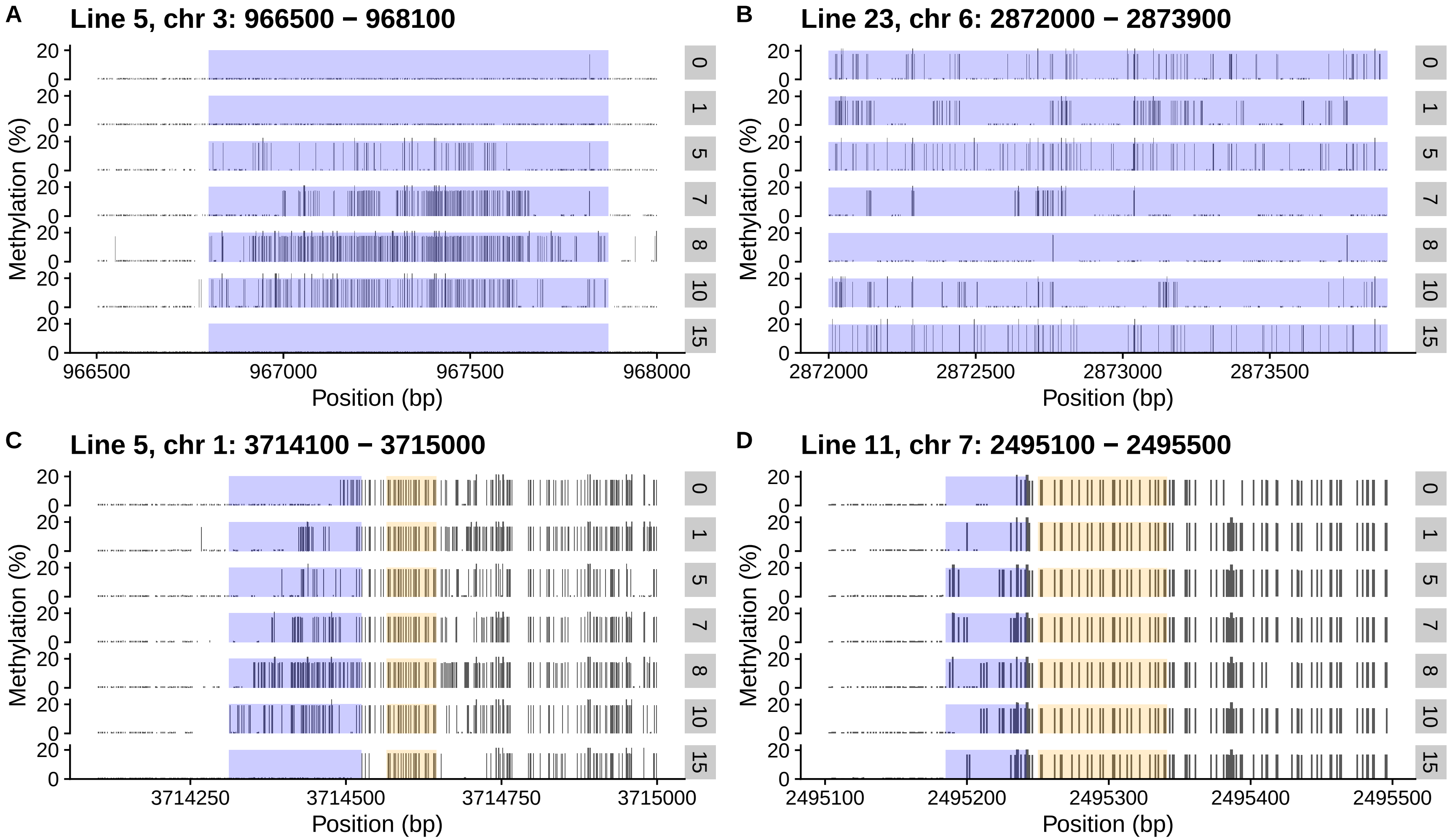

Figure S12: Examples of of loci with changes in methylation patterns in the MA lines. Tracks show methylation patterns of MA lines from different transfers for a given MA line. Track 0 is the ancestor of the corresponding MA line and 1 through 15 are the transfers assayed for DNA methylation. Regions highlighted in blue show variation in DNA methylation patterns, and regions highlighted in yellow show regions with constant methylation pattern. A) Example of a locus where an initially unmethylated region gained methylation which was subsequently lost. B) Region where methylation was lost, and then gained back later. C) Region where methylation seems to expand into an unmethylated region, while methylation downstream remains constant. D) Another region where methylation patterns change that is adjacent to region where methylation remains constant.

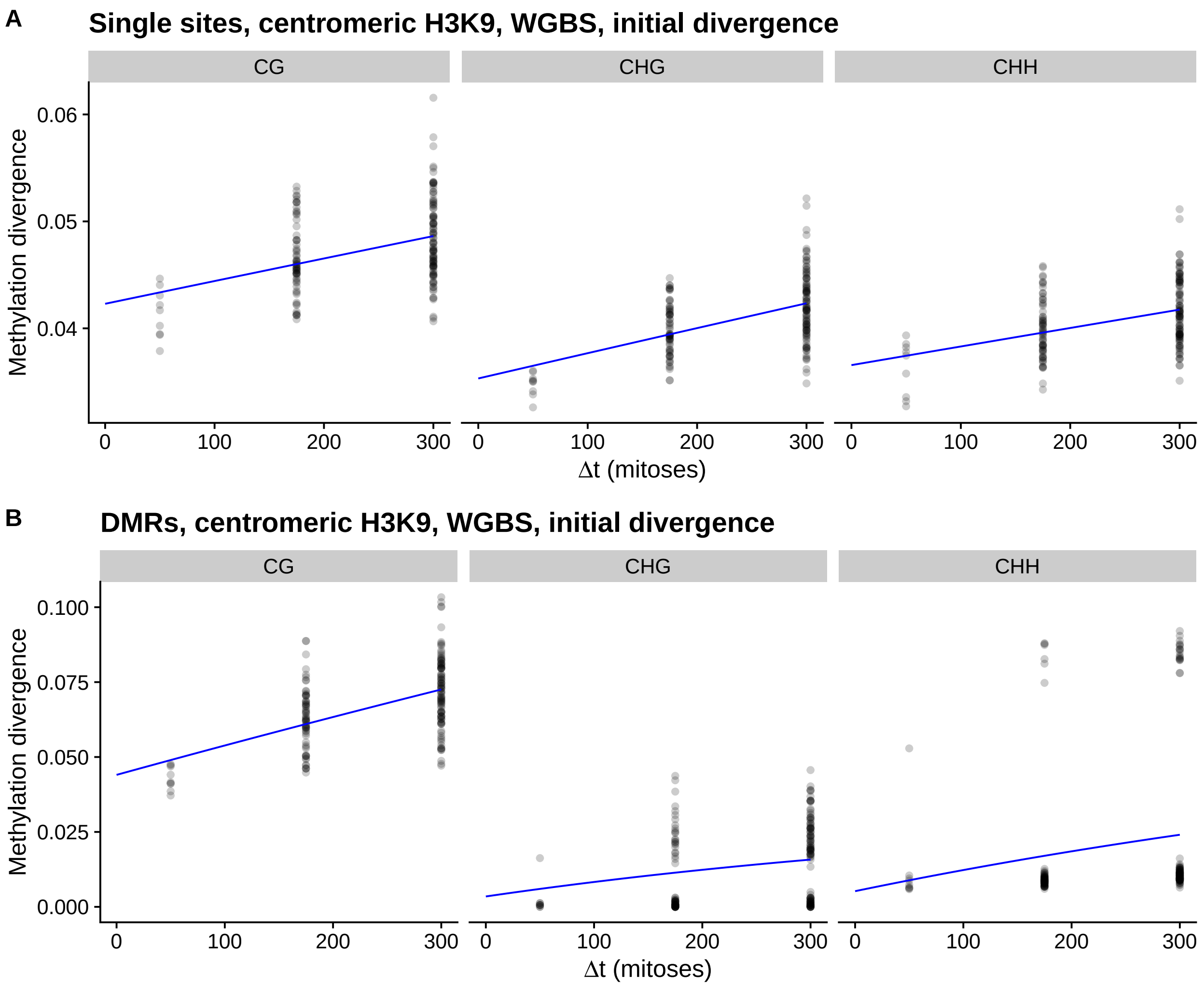

Figure S13: Pairwise divergence in methylation patterns in centromeric regions among the MA lines and their ancestors in the WGBS dataset only for the initial 300 mitoses. Blue line is a fit from the AlphaBeta neutral epimutation accumulation model. A) Divergence calculated for single sites. B) Divergence calculated for DMRs.

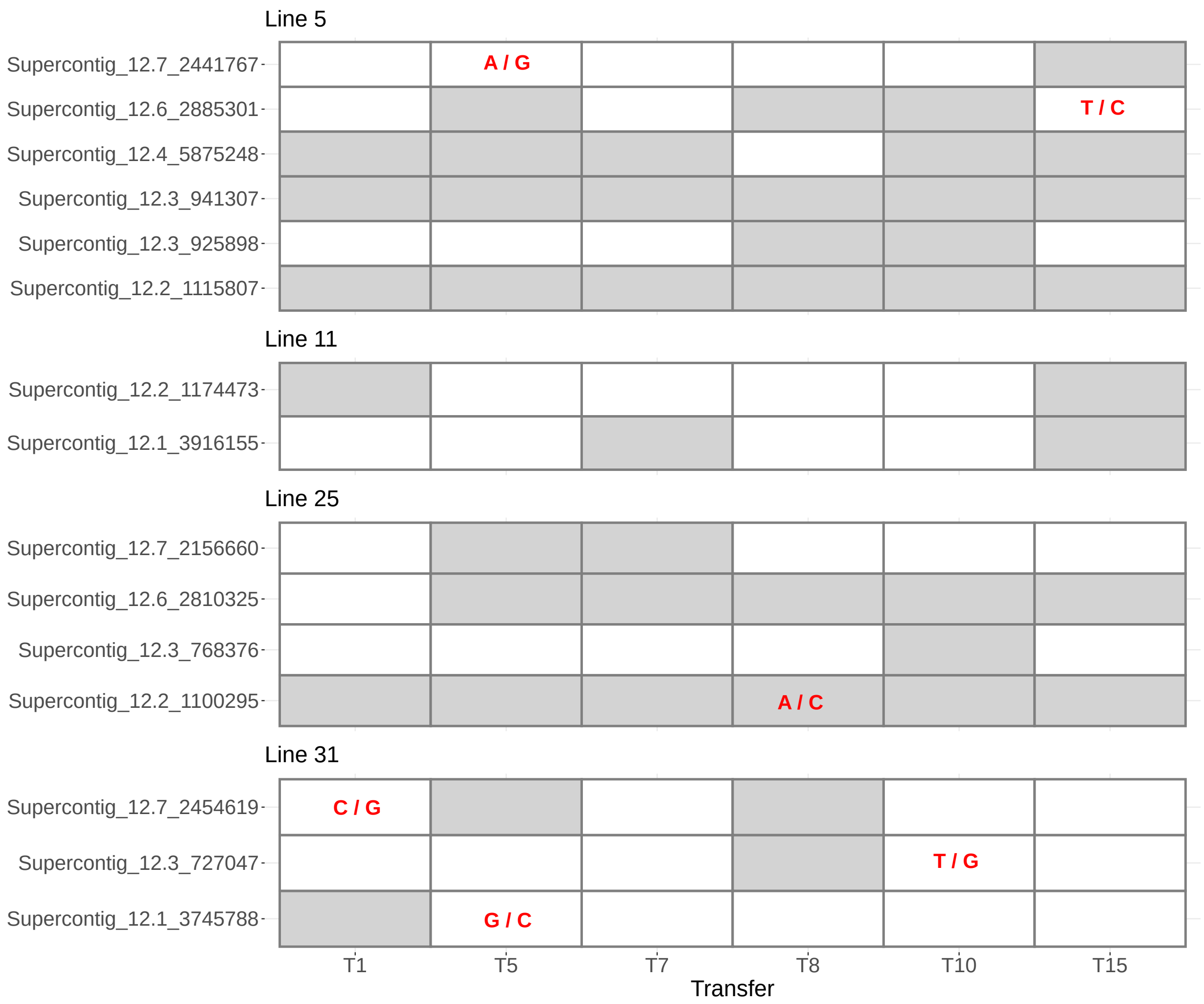

Figure S14: Differentially methylated regions (DMRs) that overlap with genetic mutations are depicted in rows and columns indicate different transfers of the MA experiment. Grey and white colours indicate different methylation states for that DMR. A genetic mutation occurring in a particular transfer is marked in red (ancestor base/derived base). Rows without genetic mutations signify DMR locations where the genetic mutation occurred after transfer fifteen, outside our Nanopore-seq dataset.

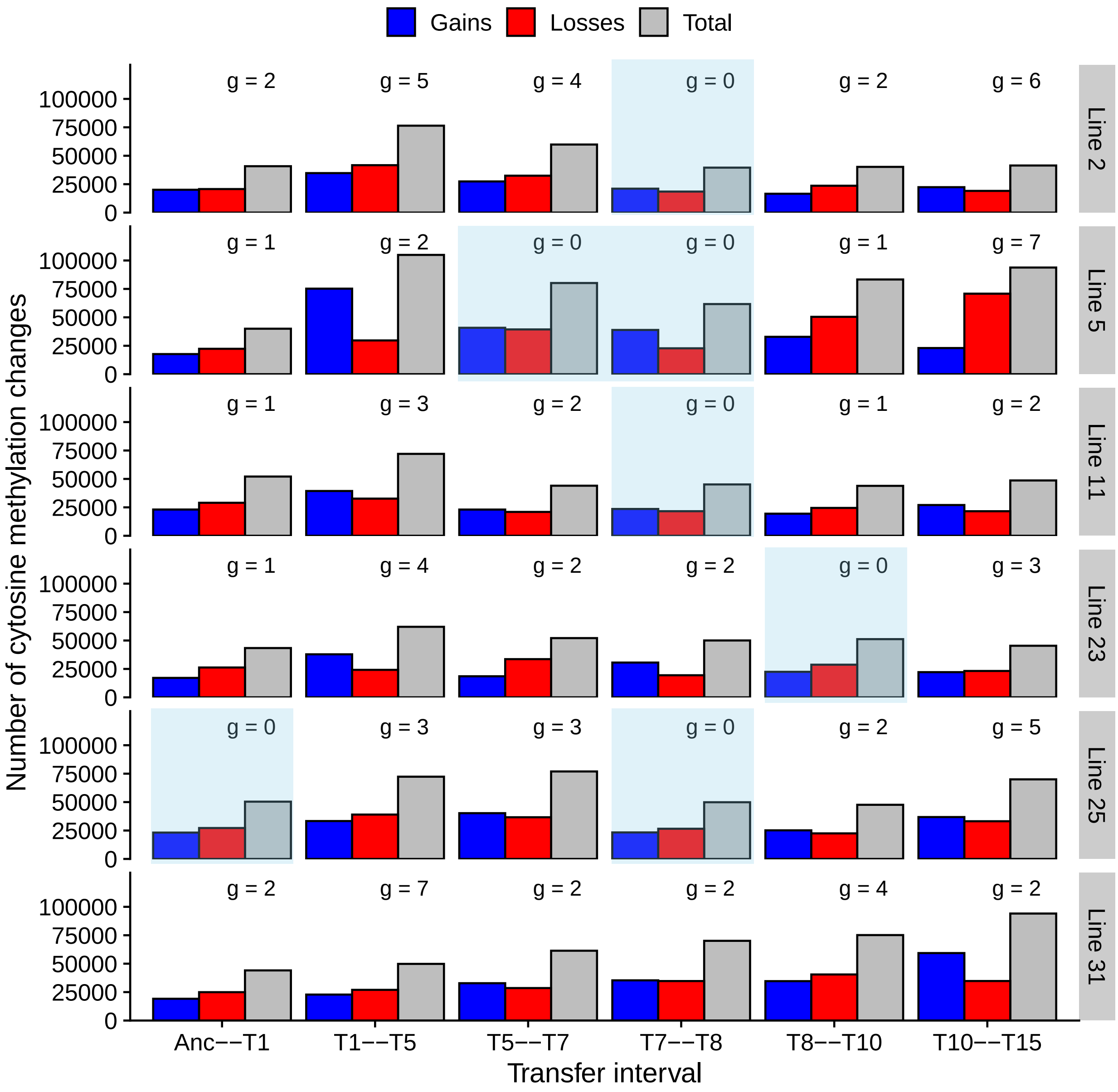

Figure S15: Number of cytosine methylation changes at each interval for the samples sequenced with Nanopore. Number of genetic mutations that happened in each interval is shown on top of the bars and intervals where no genetic mutations occurred are highlighted in blue.

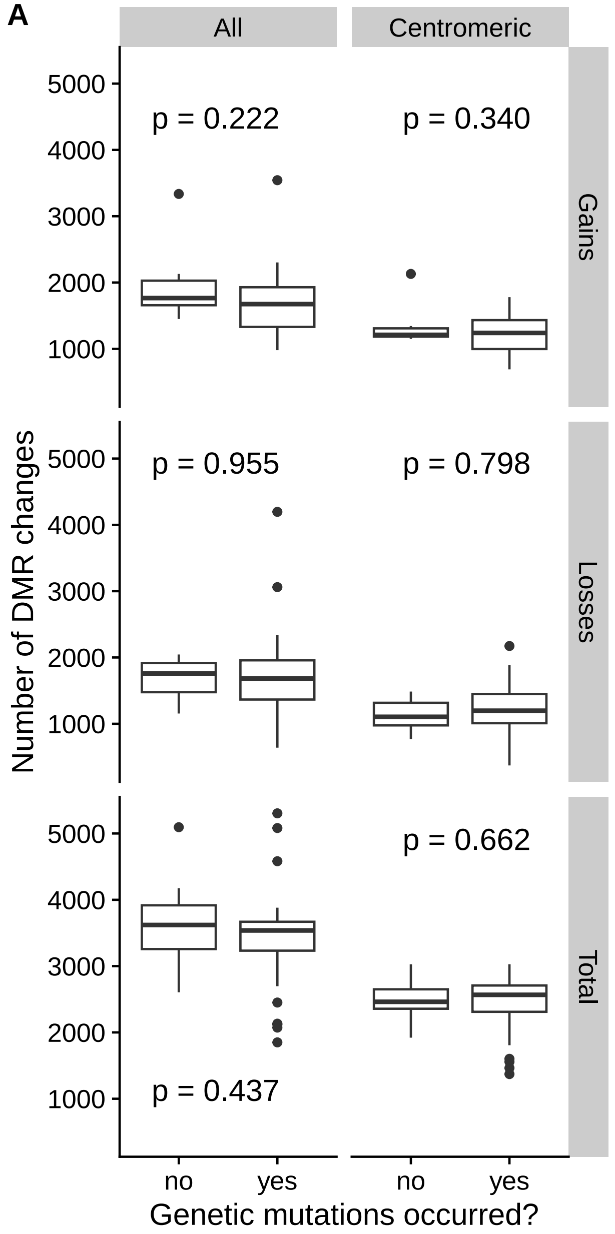

Figure S16: A) Number of DMR changes for each interval and whether genetic mutations occurred in that interval for the Nanopore-seq dataset. There were 36 intervals, and 7 intervals without genetic mutations. B) Only those intervals that contained a single transfer. There were 12 such intervals, of which five did not have a genetic mutation.

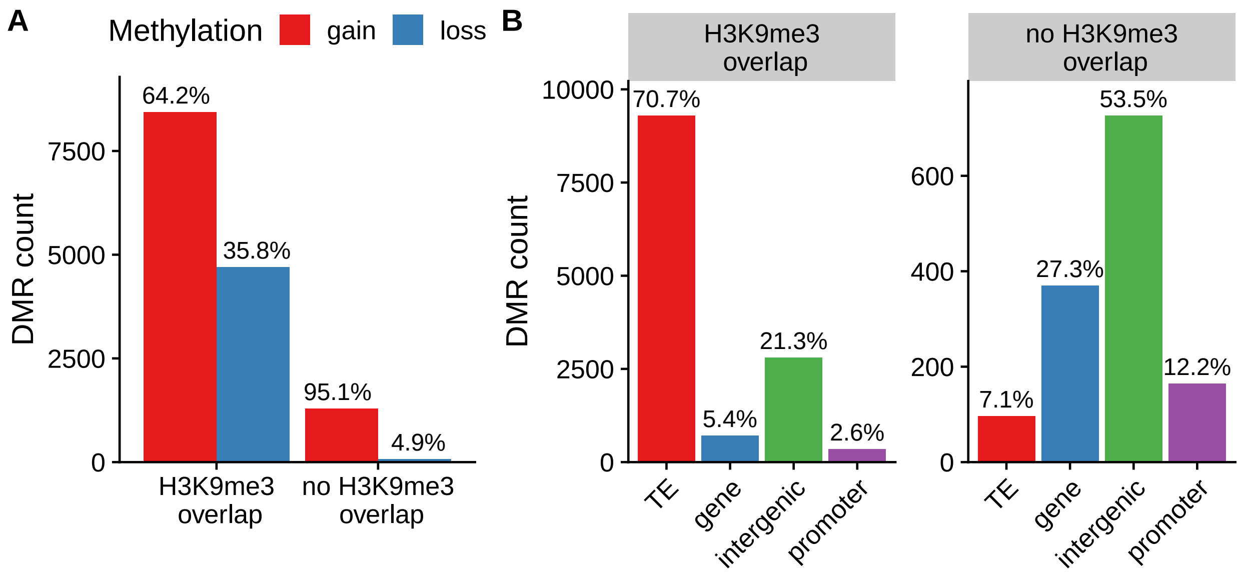

Figure S17: Properties of DMRs and whether they overlap H3K9me3. A) DMR counts and whether the DMR resulted in gain or loss of methylation for non-overlapping and for those overlapping with H3K9me3. B) Annotations of DMRs overlapping or non-overlapping H3K9me3.

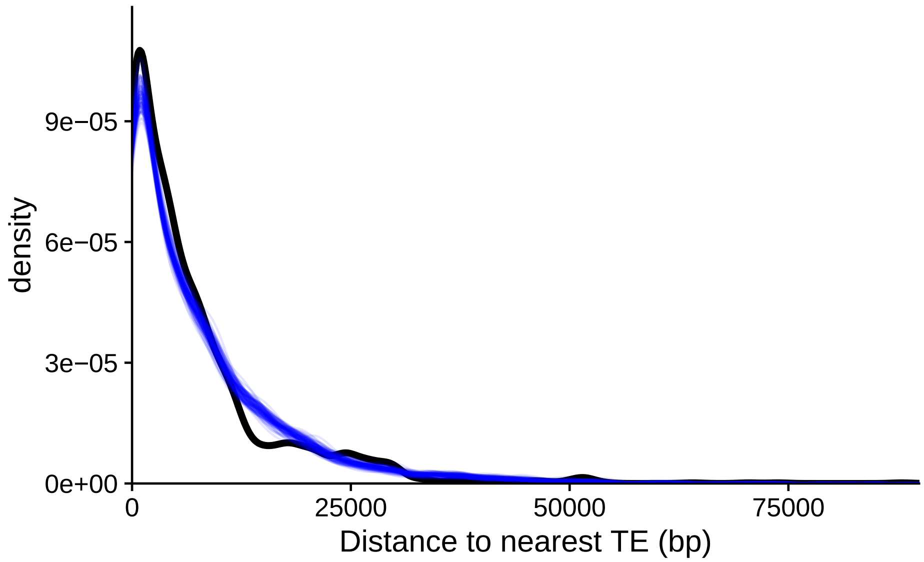

Figure S18: Black line shows the density of distances from DMRs that occurred in euchromatic genes and promoters to the nearest TE. There were 1694 DMRs in euchromatic genes, and 549 in euchromatic promoters. Blue lines represent the density of distances from random euchromatic genes and promoters to the nearest TE. One hundred independent simulations were performed, with the same number of genes and promoters sampled from euchromatin.

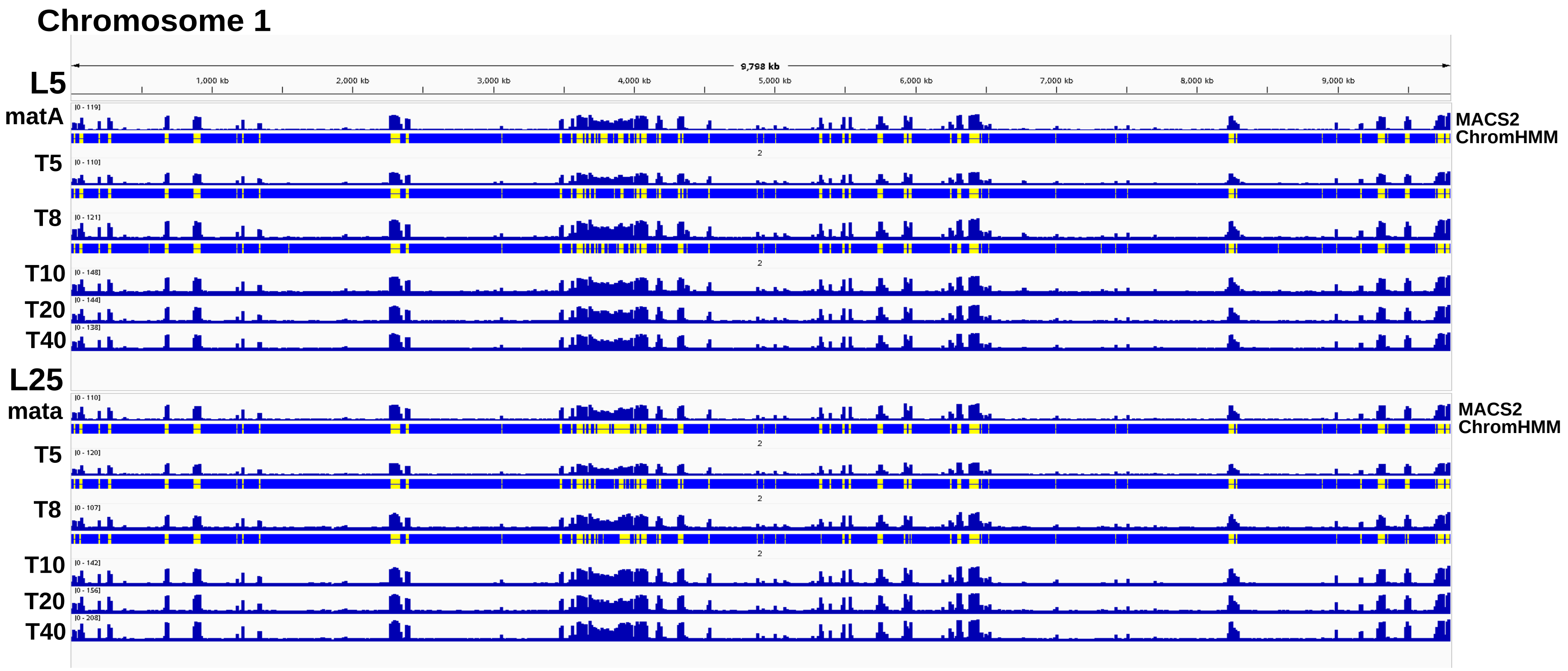

Figure S19: IGV screenshot showing lines 5 and 25 across transfers 5, 8, 10, 20, and 40. In the ancestor and the first two transfers, we include a comparison of two computational methods used to identify H3K9me3-enriched regions. Yellow fragments indicate regions predicted as H3K9me3-enriched by ChromHMM. Chromosome (Linkage Group) I is shown, as a representative example, to illustrate the absence of major changes in H3K9me3 enrichment over time.

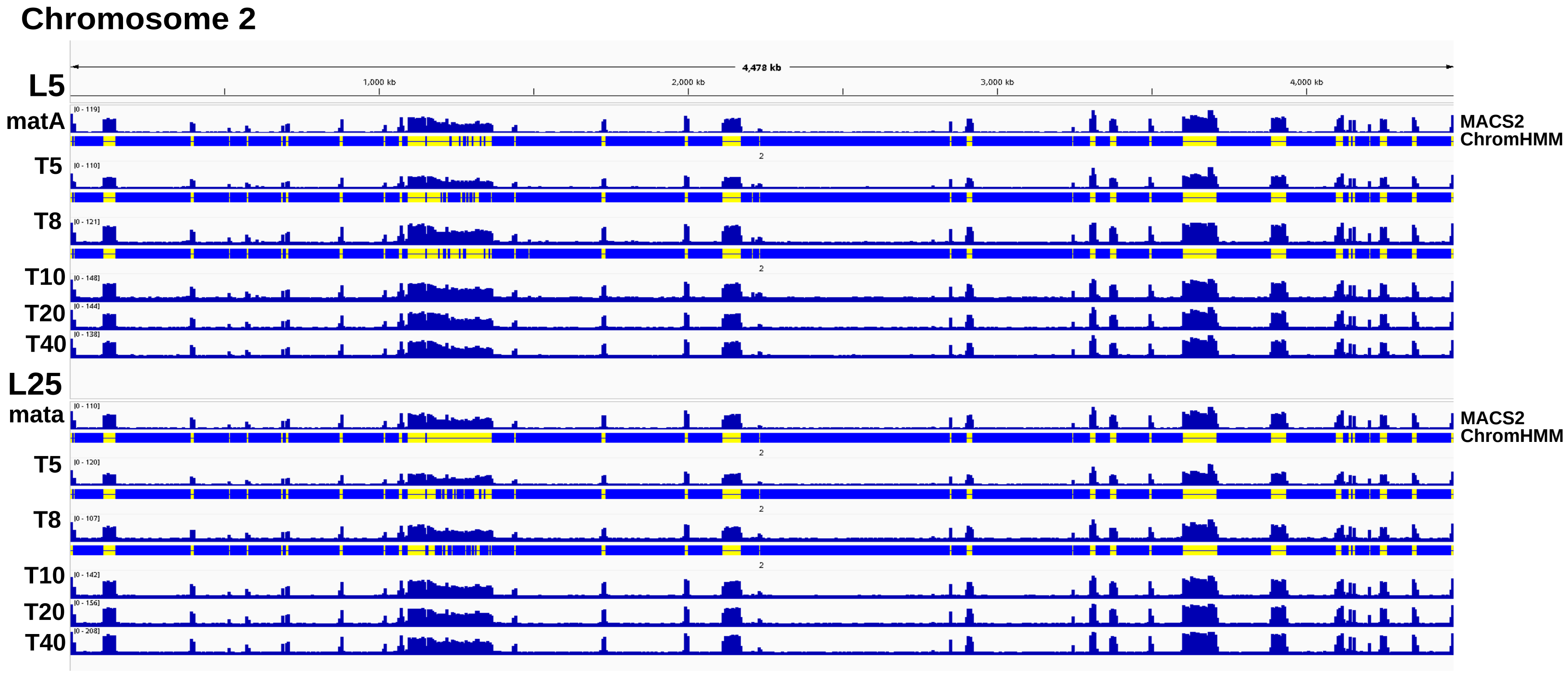

Figure S20: IGV screenshot showing lines 5 and 25 across transfers 5, 8, 10, 20, and 40. In the ancestor and the first two transfers, we include a comparison of two computational methods used to identify H3K9me3-enriched regions. Yellow fragments indicate regions predicted as H3K9me3-enriched by ChromHMM. Chromosome (Linkage Group) II is shown, as a representative example, to illustrate the absence of major changes in H3K9me3 enrichment over time.

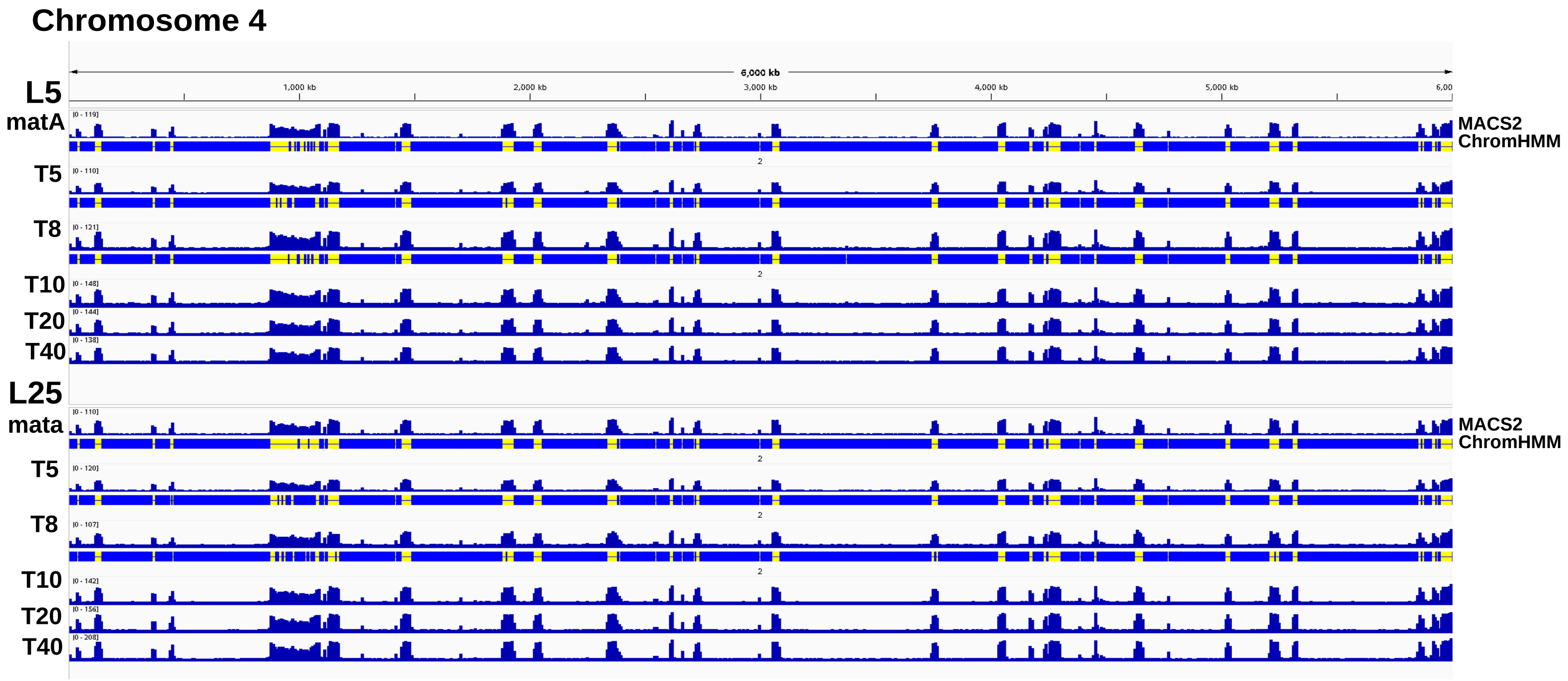

Figure S21: IGV screenshot showing lines 5 and 25 across transfers 5, 8, 10, 20, and 40. In the ancestor and the first two transfers, we include a comparison of two computational methods used to identify H3K9me3-enriched regions. Yellow fragments indicate regions predicted as H3K9me3-enriched by ChromHMM. Chromosome (Linkage Group) IV is shown, as a representative example, to illustrate the absence of major changes in H3K9me3 enrichment over time.

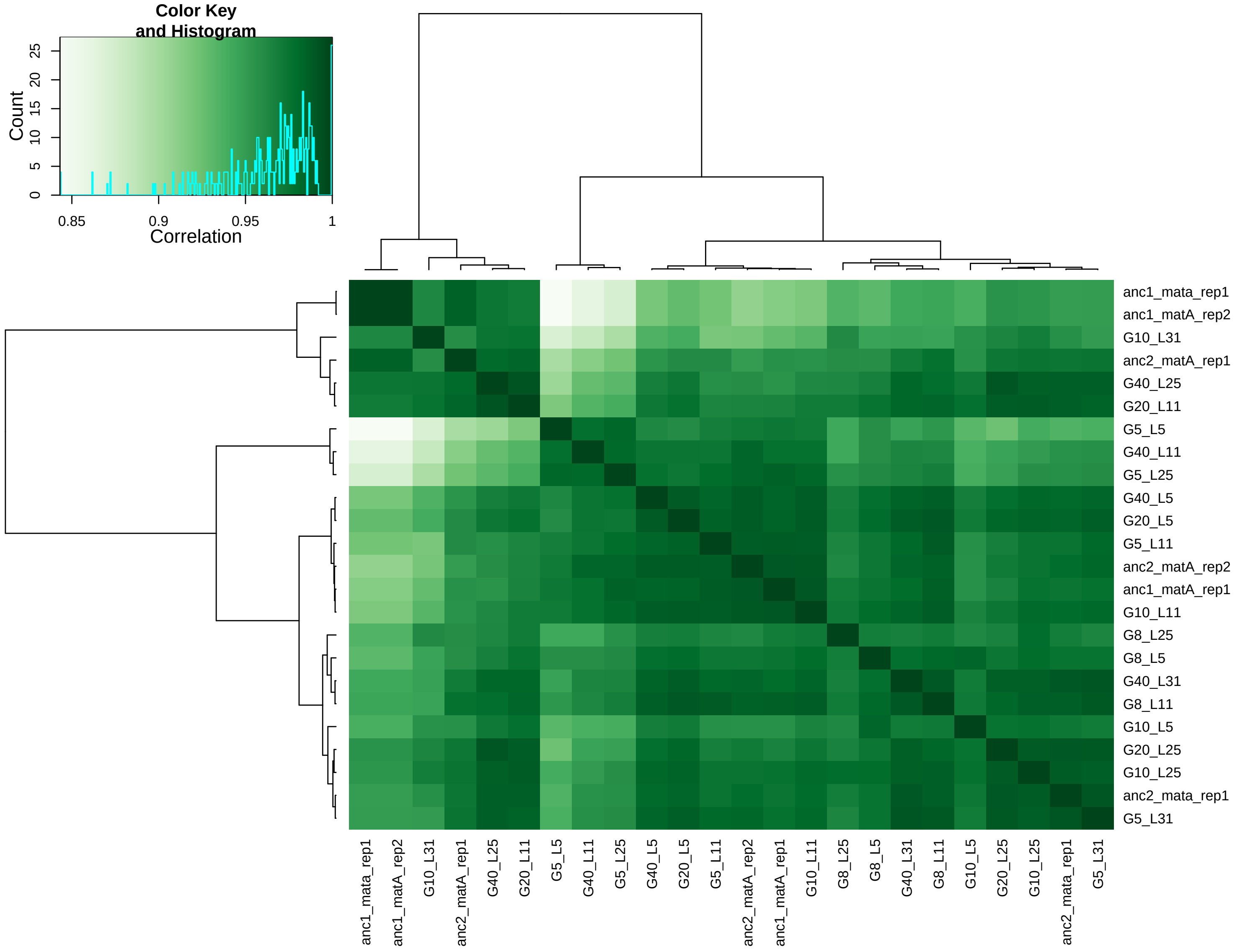

Figure S22: Correlation heatmap of H3K9me3 regions. Created with the package Diffbind. Intensities of color green represent the Pearson coefficient number

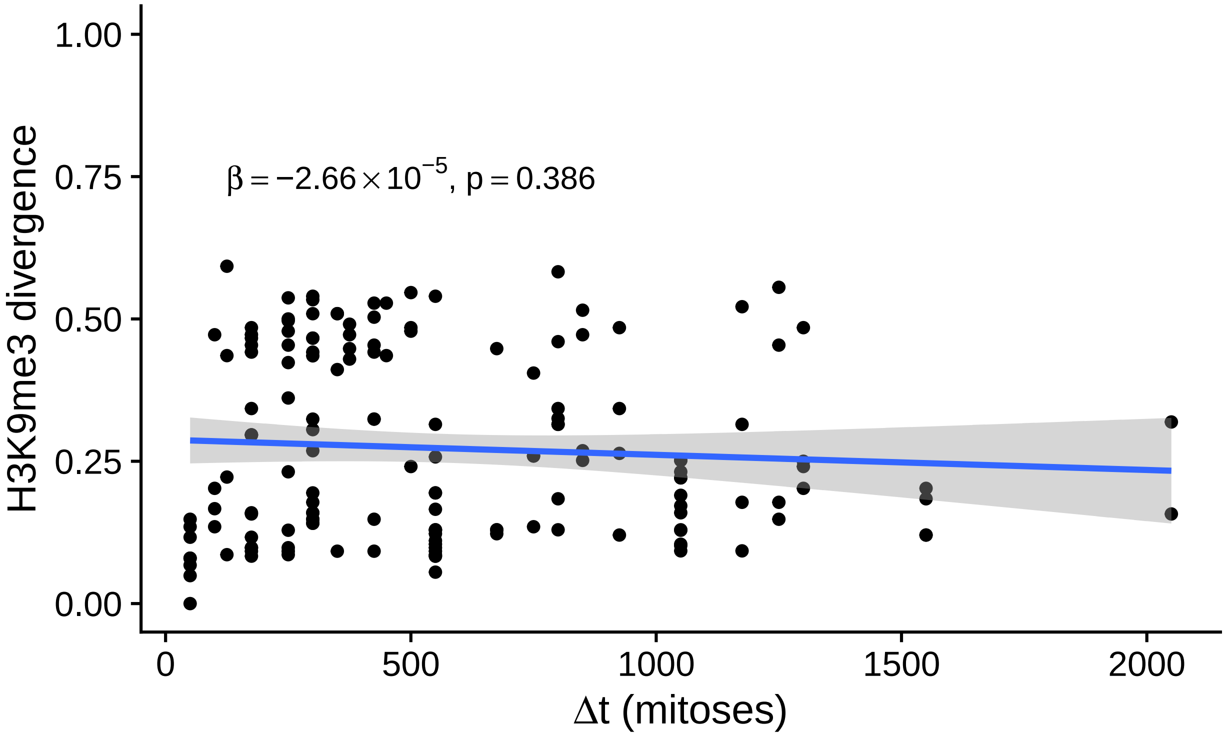

Figure S23: Divergence in presence and absence of peaks of H3K9me3 among the MA lines. The slope of the regression line of divergence in H3K9me3 against number of mitoses separating the MA lines is not statistically different from zero.

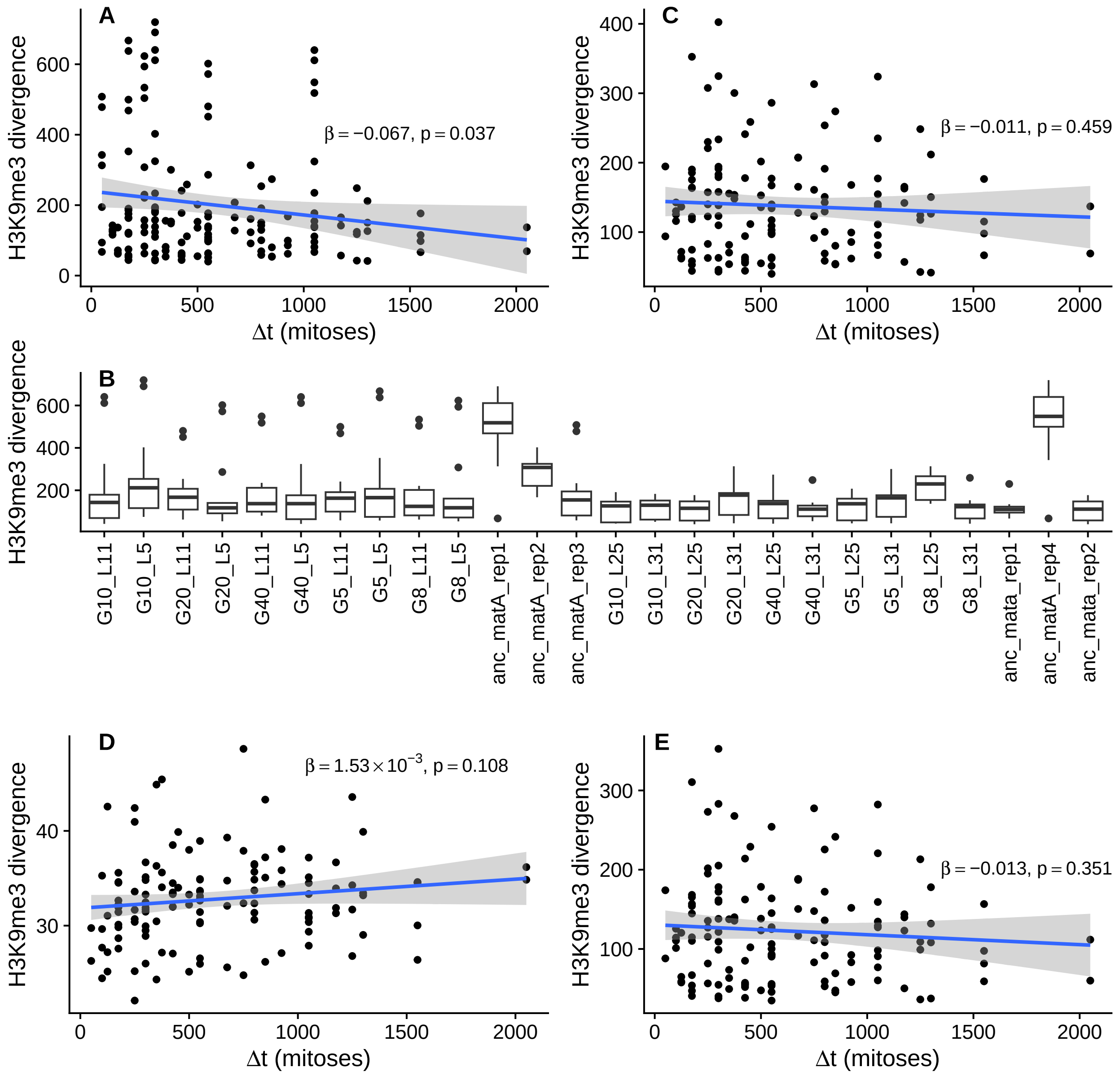

Figure S24: Divergence in H3K9me3 read counts in bins across the genome among the MA lines. A) Data for whole genome in 5kb bins, where a small negative slope was observed. B) Divergence for each sample compared to other samples within pedigree. Two samples, that are replicates of the mat A ancestor, have considerably higher divergence to other samples. C) Reanalysis of divergence across the whole genome, with the two outlier samples detected in B) excluded. Divergence does not change significantly as the MA lines diverge. D) Data for centromeric regions only in 500 bp bins, with two outlier samples are excluded; the slope of divergence is not significantly different from zero. E) Data for regions excluding the centromeres in 5 kb bins, with two outlier samples excluded; the slope of divergence is not significantly different from zero.

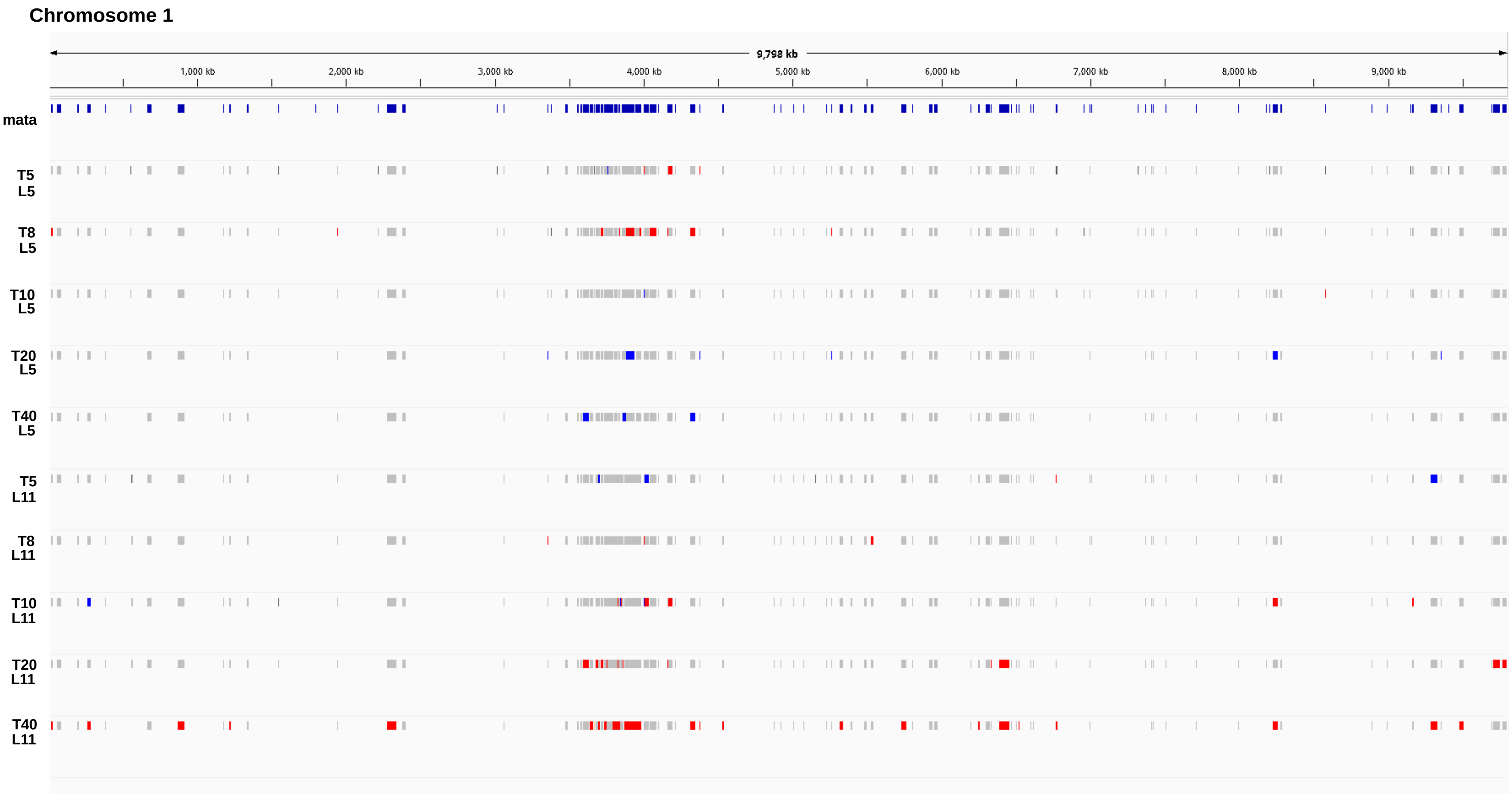

Figure S25: Results of ChromTime analysis. Representative examples showing lines 5 and 11, limited to chromosome 1. Grey regions indicate steady H3K9me3 domains, red denotes regions classified as expanded, and blue indicates regions classified as contracted over time.

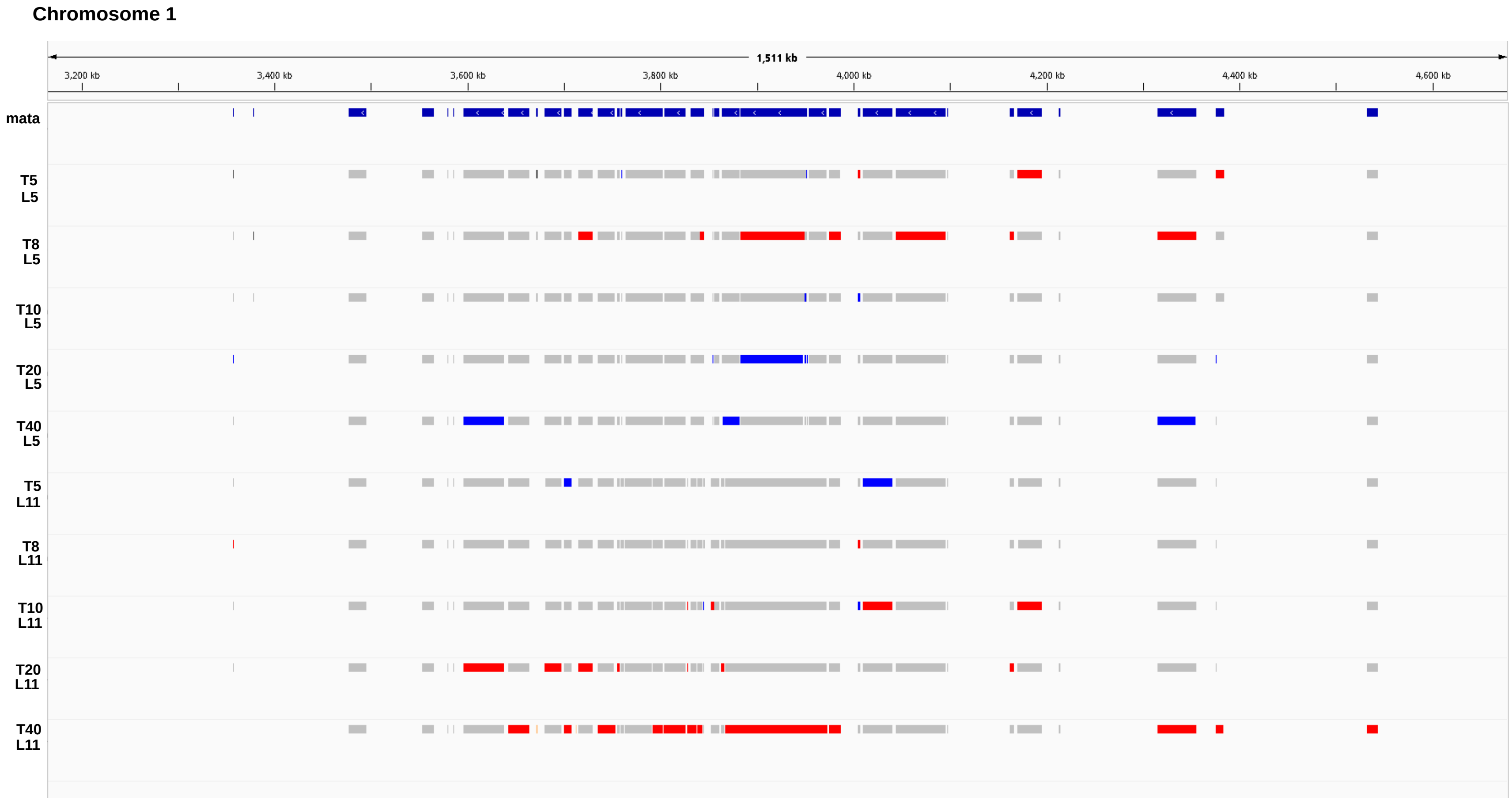

Figure S26: Results of ChromTime analysis. Representative examples showing lines 5 and 11, limited to centromeric regions of chromosome 1. Grey regions indicate steady H3K9me3 domains, red denotes regions classified as expanded, and blue indicates regions classified as contracted over time. Zoom-in on the centromeric region shows that many of the regions labeled as expansions or contractions appear to be technical artifacts, rather than true biological changes.

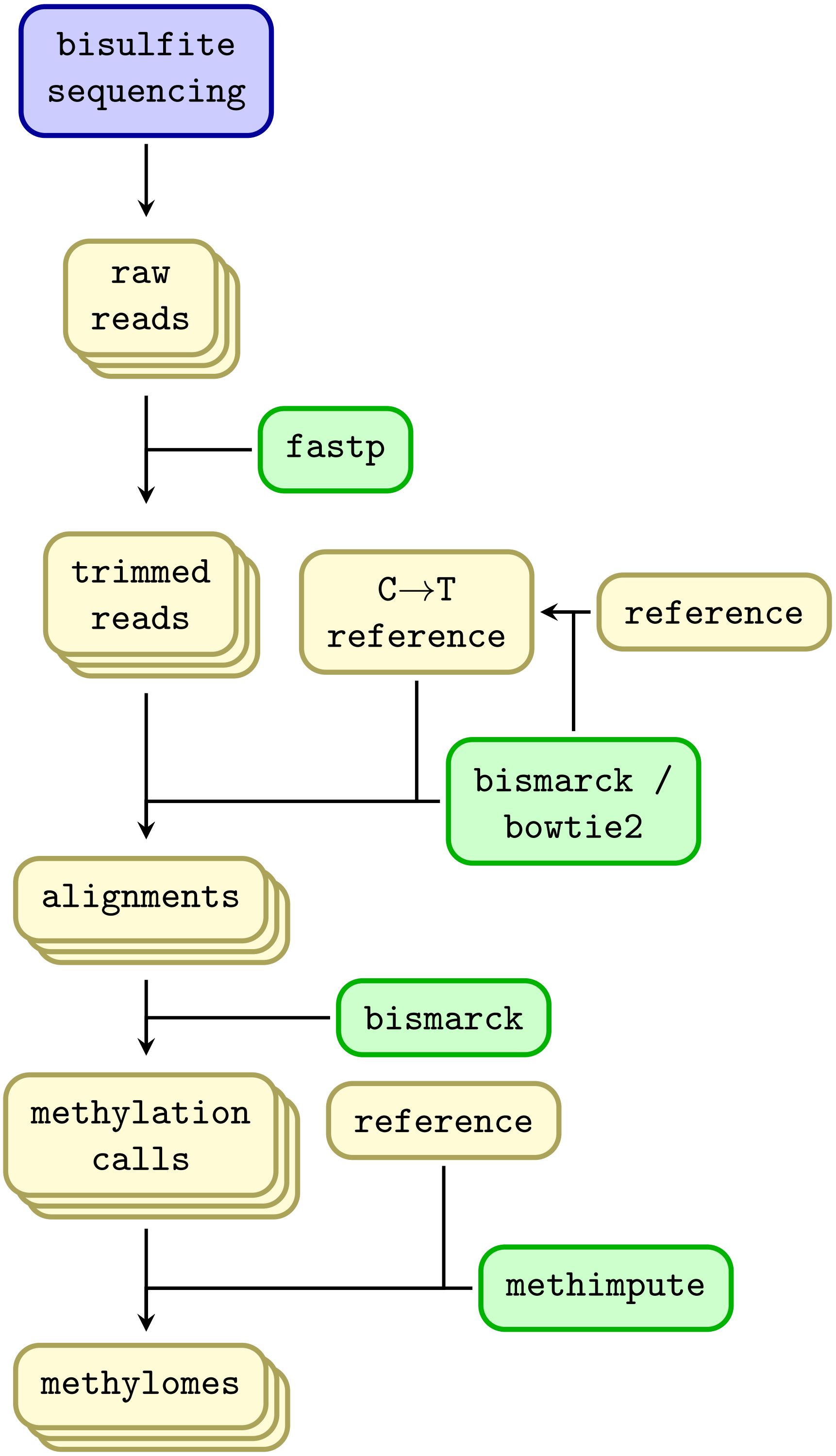

Figure S27: An overview of the bioinformatics workflow for processing WGBS samples. Yellow boxes are files, and green boxes represent programs.

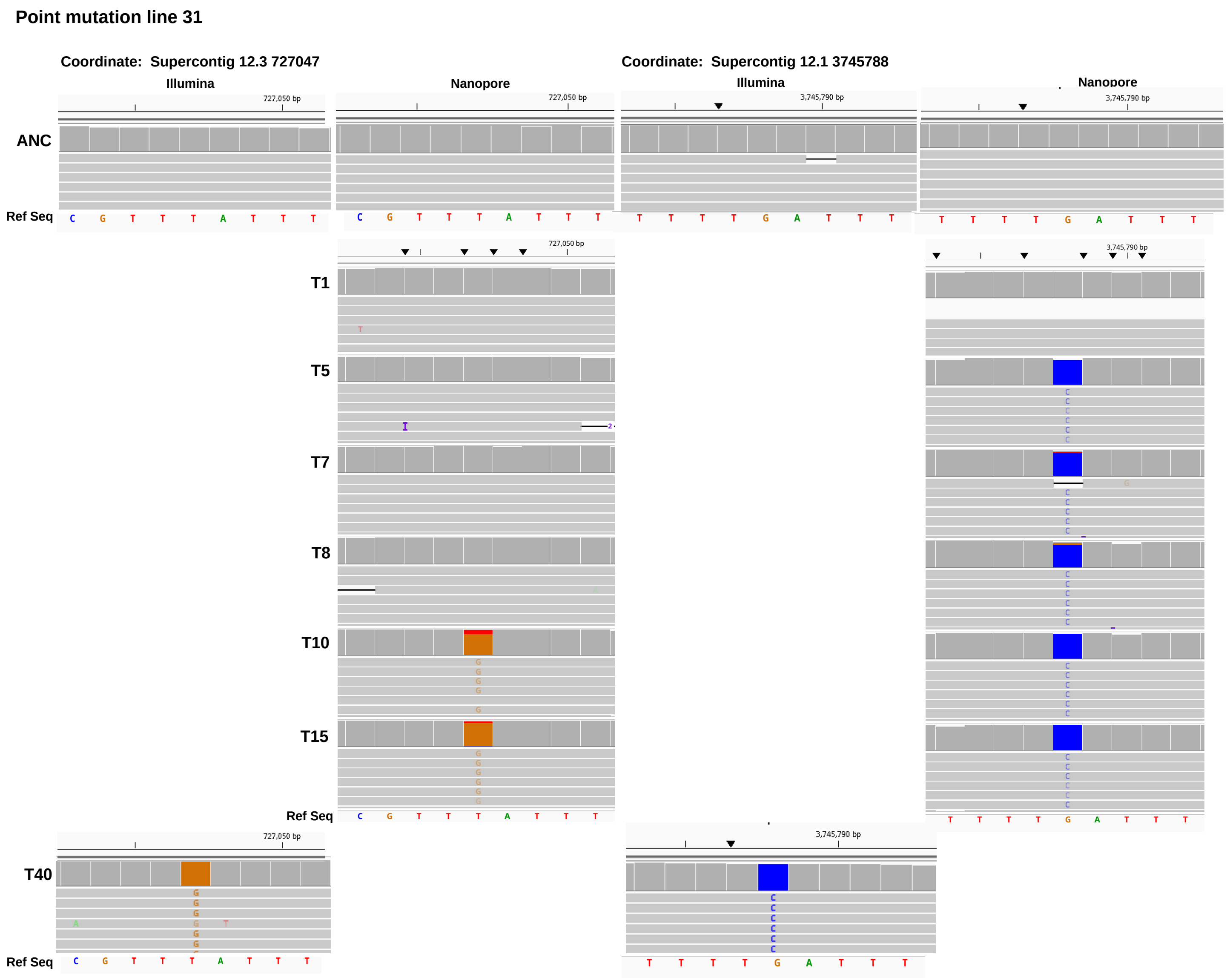

Figure S28: IGV screenshots showing single-nucleotide polymorphisms (SNPs) identified in MA line 31. Shown are representative visualizations of point mutations detected by Nanopore and Illumina sequencing: a T→G substitution in transfer 10 at chromosome 3, position 717,047, and a G→C substitution in transfer 5 at chromosome 1, position 3,745,788. For each mutation, we present screenshots of the ancestor sequenced by Illumina and Nanopore at the mutation coordinate, followed by intermediate transfers sequenced by Nanopore and transfer 40 sequenced by Illumina.

#### 1.4 Supplementary Tables

Table S1: Samples sequenced with bisulfite sequencing. Lines 1 through 20 are descended from the *mat A* ancestor, and lines 21 through 40 from the *mat a* ancestor. Bs. eff. is bisulfite conversion efficiency estimated from $\lambda$-phage spike-in control.

| Transfer | Line | Reads | Depth (x) | Coverage (%) | Bs. eff. (%) |
|---|---|---|---|---|---|
| Ancestor | *mat A* rep 1 | 15129280 | 53 | 99.83 | 99.72 |
| Ancestor | *mat A* rep 2 | 14203858 | 50 | 99.86 | 99.73 |
| Ancestor | *mat A* rep 3 | 11905318 | 41 | 99.85 | 99.72 |
| Ancestor | *mat a* rep 1 | 12009736 | 42 | 99.80 | 99.72 |
| Ancestor | *mat a* rep 2 | 13310900 | 46 | 99.83 | 99.76 |
| Ancestor | *mat a* rep 3 | 12958022 | 45 | 99.83 | 99.75 |
| 5 | L1 | 13352788 | 47 | 99.86 | 99.73 |
| 5 | L6 | 13224652 | 46 | 99.85 | 99.76 |
| 5 | L7 | 12854380 | 45 | 99.84 | 99.76 |
| 5 | L10 rep 1 | 14067628 | 49 | 99.85 | 99.74 |
| 5 | L10 rep 2 | 14652934 | 51 | 99.86 | 99.75 |
| 5 | L13 | 14044374 | 49 | 99.85 | 99.75 |
| 5 | L14 | 14065732 | 49 | 99.85 | 99.76 |
| 5 | L15 | 13833630 | 48 | 99.84 | 99.73 |
| 5 | L17 | 13354348 | 47 | 99.83 | 99.73 |
| 5 | L19 | 11498660 | 40 | 99.83 | 99.75 |
| 5 | L20 | 14094228 | 49 | 99.82 | 99.76 |
| 5 | L21 | 13514998 | 47 | 99.81 | 99.74 |
| 5 | L28 | 13554304 | 48 | 99.84 | 99.74 |
| 5 | L29 | 12014028 | 42 | 99.80 | 99.75 |
| 5 | L30 | 12645992 | 44 | 99.77 | 99.76 |
| 5 | L33 | 14542396 | 51 | 99.82 | 99.74 |
| 5 | L34 | 15461790 | 54 | 99.83 | 99.72 |
| 5 | L35 | 13356940 | 47 | 99.81 | 99.71 |
| 5 | L36 | 9685796 | 34 | 99.82 | 99.75 |
| 5 | L39 | 14899702 | 52 | 99.83 | 99.75 |
| 5 | L40 | 15231878 | 53 | 99.83 | 99.74 |
| 20 | L1 | 16455520 | 58 | 99.81 | 99.79 |
| 20 | L6 | 16536660 | 58 | 99.85 | 99.78 |
| 20 | L7 | 16499612 | 58 | 99.84 | 99.77 |
| 20 | L10 rep 1 | 13837376 | 48 | 99.84 | 99.76 |
| 20 | L10 rep 2 | 12983524 | 45 | 99.86 | 99.72 |
| 20 | L13 | 13001046 | 45 | 99.86 | 99.73 |

Continued on next page...

Table S1 – continued from previous page

| Transfer | Line | Reads | Depth (x) | Coverage (%) | Bs. eff. (%) |
|---|---|---|---|---|---|
| 20 | L14 | 13140234 | 46 | 99.84 | 99.75 |
| 20 | L15 | 1396670 | 49 | 99.84 | 99.76 |
| 20 | L17 | 18664836 | 65 | 99.85 | 99.76 |
| 20 | L19 | 11421242 | 40 | 99.83 | 99.74 |
| 20 | L20 | 14105642 | 49 | 99.72 | 99.76 |
| 20 | L21 | 17244100 | 61 | 99.32 | 99.76 |
| 20 | L28 | 16031346 | 56 | 99.80 | 99.75 |
| 20 | L29 | 19810272 | 69 | 99.84 | 99.75 |
| 20 | L30 | 16756842 | 59 | 99.79 | 99.77 |
| 20 | L33 | 16477048 | 58 | 99.81 | 99.75 |
| 20 | L34 | 13845756 | 48 | 99.82 | 99.75 |
| 20 | L35 | 18711334 | 66 | 99.81 | 99.75 |
| 20 | L36 | 18650916 | 65 | 99.82 | 99.75 |
| 20 | L39 | 12331098 | 43 | 99.82 | 99.77 |
| 20 | L40 | 12583648 | 44 | 99.82 | 99.75 |
| 40 | L1 | 14543480 | 51 | 99.83 | 99.75 |
| 40 | L6 | 15174730 | 53 | 99.66 | 99.69 |
| 40 | L7 | 12888682 | 45 | 99.80 | 99.70 |
| 40 | L10 rep 1 | 12976256 | 45 | 99.82 | 99.69 |
| 40 | L10 rep 2 | 13621346 | 48 | 99.86 | 99.73 |
| 40 | L13 | 15574080 | 54 | 99.82 | 99.67 |
| 40 | L14 | 14154158 | 49 | 99.83 | 99.69 |
| 40 | L15 | 16945084 | 59 | 99.83 | 99.72 |
| 40 | L17 | 16308294 | 57 | 99.84 | 99.68 |
| 40 | L19 | 14457268 | 50 | 99.82 | 99.64 |
| 40 | L20 | 18716914 | 66 | 99.72 | 99.67 |
| 40 | L21 | 15669808 | 55 | 99.18 | 99.69 |
| 40 | L28 | 17091676 | 60 | 99.80 | 99.65 |
| 40 | L29 | 14925692 | 52 | 99.81 | 99.63 |
| 40 | L30 | 13043906 | 46 | 99.81 | 99.67 |
| 40 | L33 | 12954644 | 45 | 99.77 | 99.22 |
| 40 | L34 | 17665520 | 62 | 99.82 | 99.67 |
| 40 | L35 | 13437214 | 47 | 99.77 | 99.73 |
| 40 | L36 | 17087492 | 60 | 99.80 | 99.69 |
| 40 | L39 | 13776582 | 48 | 99.81 | 99.72 |
| 40 | L40 | 18100634 | 63 | 99.83 | 99.71 |

Table S2: Samples sequenced with Nanopore sequencing. Length is the median Nanopore read length. Lines 2, 5, and 11 are descended from the *mat A* ancestor, and lines 23, 25, and 31 from the *mat a* ancestor.

| Transfer | Line | Reads | Length | Coverage (%) | Depth (x) |
|---|---|---|---|---|---|
| Ancestor | *mat A* | 226810 | 3554.5 | 99.97 | 37.05 |
| Ancestor | *mat a* | 307827 | 3302 | 99.97 | 43.69 |
| 1 | L2 | 355489 | 2576 | 99.97 | 31.63 |
| 1 | L5 | 355210 | 2844 | 99.97 | 34.13 |
| 1 | L11 | 470989 | 2971 | 99.97 | 47.59 |
| 1 | L23 | 368655 | 3496 | 99.97 | 42.92 |
| 1 | L25 | 301674 | 2782 | 99.97 | 27.49 |
| 1 | L31 | 318778 | 3799 | 99.97 | 42.48 |
| 5 | L2 | 588728 | 2905 | 99.97 | 58.44 |
| 5 | L11 | 260810 | 3133 | 99.97 | 26.91 |
| 5 | L5 | 532908 | 3081 | 99.97 | 57.96 |
| 5 | L23 | 290791 | 2667 | 99.97 | 26.16 |
| 5 | L25 | 450272 | 3306 | 99.96 | 51.42 |
| 5 | L31 | 266086 | 3535 | 99.97 | 31.67 |
| 7 | L2 | 326525 | 3357 | 99.97 | 37.42 |
| 7 | L5 | 207747 | 4098 | 99.97 | 34.9 |
| 7 | L11 | 233683 | 3283 | 99.97 | 27.29 |
| 7 | L23 | 220030 | 4507 | 99.97 | 34.41 |
| 7 | L25 | 204732 | 3552 | 99.96 | 25.99 |
| 7 | L31 | 391108 | 3551 | 99.97 | 52.36 |
| 8 | L2 | 214994 | 3982 | 99.97 | 34.06 |
| 8 | L5 | 241171 | 3514 | 99.97 | 33.86 |
| 8 | L11 | 215031 | 4516 | 99.97 | 36.36 |
| 8 | L23 | 220595 | 3474 | 99.96 | 27.22 |
| 8 | L25 | 262599 | 3653 | 99.96 | 38.25 |
| 8 | L31 | 232717 | 3549 | 99.96 | 30.15 |
| 10 | L2 | 223022 | 4201 | 99.97 | 34.85 |
| 10 | L5 | 399996 | 3563 | 99.98 | 53.49 |
| 10 | L11 | 197851 | 4244 | 99.97 | 33.29 |
| 10 | L23 | 322348 | 3427 | 99.96 | 39.28 |
| 10 | L25 | 140407 | 5228 | 99.96 | 28.22 |
| 10 | L31 | 371921 | 3575 | 99.96 | 53.51 |
| 15 | L2 | 244290 | 3561 | 99.97 | 33.45 |
| 15 | L5 | 251497 | 3724 | 99.96 | 37.62 |
| 15 | L11 | 203485 | 3774 | 99.97 | 28.9 |
| 15 | L23 | 225741 | 3723 | 99.97 | 33.03 |
| 15 | L25 | 399885 | 3511 | 99.98 | 53.32 |
| 15 | L31 | 167378 | 3536 | 99.95 | 25.6 |

Table S3: Samples sequenced with ChIP-sequencing with antibody against H3K9me3. Lines 5 and 11 are descened from the *mat A* ancestor, and lines 25 and 31 from the *mat a* ancestor.

| Transfer | Line | Reads | Depth (x) |
|---|---|---|---|
| Ancestor | *mat a* rep 1 | 4406326 | 13 |
| Ancestor | *mat a* rep 2 | 8573262 | 25 |
| Ancestor | *mat A* rep 1 | 16456278 | 48 |
| Ancestor | *mat A* rep 2 | 3734610 | 11 |
| Ancestor | *mat A* rep 3 | 4477949 | 13 |
| Ancestor | *mat A* rep 4 | 7454673 | 22 |
| 5 | L5 | 3860381 | 11 |
| 5 | L11 | 4643805 | 13 |
| 5 | L25 | 3833040 | 11 |
| 5 | L31 | 7346841 | 21 |
| 8 | L5 | 6753961 | 20 |
| 8 | L11 | 6179668 | 18 |
| 8 | L25 | 6496179 | 19 |
| 8 | L31 | 21930090 | 64 |
| 10 | L5 | 9824968 | 29 |
| 10 | L11 | 5656063 | 16 |
| 10 | L25 | 7835358 | 23 |
| 10 | L31 | 1077318 | 32 |
| 20 | L5 | 6510277 | 19 |
| 20 | L11 | 9720105 | 29 |
| 20 | L25 | 10277927 | 30 |
| 20 | L31 | 23942748 | 71 |
| 40 | L5 | 6019916 | 18 |
| 40 | L11 | 3935274 | 11 |
| 40 | L25 | 11786931 | 35 |
| 40 | L31 | 6697276 | 20 |
